# Ribosomal protein control of hematopoietic stem cell transformation through regulation of metabolism

**DOI:** 10.1101/2023.05.31.543132

**Authors:** Bryan Harris, Dinesh K. Singh, Billy Truong, Michele Rhodes, Rachael Price, Susan Shinton, Monika Verma, Bridget Aylward, Shawn P. Fahl, Shanna R. Sprinkle, Sarah Aminov, Minshi Wang, Yong Zhang, Jaqueline Perrigoue, Rachel Kessel, Suraj Peri, Joshua West, Orsi Giricz, Jacqueline Boultwood, Andrea Pellagatti, KH Ramesh, Cristina Montagna, Kith Pradhan, Jeffrey W. Tyner, Brian K. Kennedy, Michael Holinstat, Ulrich Steidl, Stephen Sykes, Amit Verma, David L. Wiest

## Abstract

We report here that expression of the ribosomal protein, RPL22, is frequently reduced in human myelodysplastic syndrome (MDS) and acute myelogenous leukemia (AML); and, reduced RPL22 expression is associated with worse outcomes. Mice null for Rpl22 display characteristics of an MDS-like syndrome and develop leukemia at an accelerated rate. Rpl22-deficient mice also display enhanced hematopoietic stem cell (HSC) self-renewal and obstructed differentiation potential, which arises not from reduced protein synthesis but from altered metabolism including increased fatty acid oxidation (FAO) and a striking induction of the stemness factor Lin28b in the resulting leukemia. Lin28b promotes a substantial increase in lipid content, upon which the survival of Rpl22-deficient leukemias depends. Altogether, these findings reveal that Rpl22 insufficiency enhances the leukemia potential of HSC through regulation of FAO and promotes leukemogenesis through Lin28b promotion of lipid synthesis.

**Highlights:**

- RPL22 insufficiency is observed in MDS/AML and is associated with reduced survival
- Rpl22-deficiency produces an MDS-like syndrome and facilitates leukemogenesis
- Rpl22-deficiency does not impair global protein synthesis by HSC
- Rpl22 controls leukemia survival through control of lipid synthesis

**eTOC:** Rpl22 controls the function and transformation potential of hematopoietic stem cells through regulation of lipid metabolism.

## INTRODUCTION

Ribosomopathies are a group of inherited diseases resulting from mutations or deletions of ribosomal protein (RP) encoding genes or factors that facilitate ribosome biogenesis ^1–3^. This class of diseases is characterized by defects in hematopoiesis that often culminate in complete bone marrow failure, with many patients progressing to the development of myelodysplastic syndromes (MDS) and acute myelogenous leukemia (AML) ^4^, as a result of the aberrant function of hematopoietic stem cells (HSC) ^5^. The association between the inactivation of RP and increased risk for myeloid malignancy has been known for some time, but the mechanistic link remains unclear and controversial ^6^. Some evidence suggests that RP mutations disrupt hematopoiesis and increase cancer risk by generalized impairment of ribosome function ^7–9^, while others suggest that particular RNA-binding RP are capable of performing specific regulatory functions ^10–12^. Indeed, *RPS14* haploinsufficiency is reported to promote the 5q- subtype of MDS through S100A8-mediated induction of p53 ^13^; however, the mechanistic link for this effect remains to be established. Several other studies have highlighted “extraribosomal”, regulatory functions of RP involving their ability to bind to RNA targets and modulate processes, including pre-mRNA splicing and translation ^12,14,15^. These findings highlight a need to systematically investigate the contribution of RP inactivation, or reductions in their expression, to the etiology of myeloid neoplasms to gain insight into how the lost regulatory functions of RP might contribute to malignant hematopoiesis.

One RP of particular interest is Rpl22, an RNA-binding RP that binds to the 28S rRNA and is located on its exterior surface of the 60S ribosomal subunit ^16,17^. While this highly conserved RP is ubiquitously expressed in mammalian tissues, it is distinguished from many other RP in that it is not required for ribosome biogenesis or global protein translation ^18^. Moreover, the germline ablation of the *Rpl22* gene in mice does not result in lethality or gross developmental abnormalities ^19^. Rather, Rpl22-deficiency causes selective alterations in lymphopoiesis, consistent with the loss of a regulatory function on which selected cell types are particularly dependent. Indeed, Rpl22-deficiency attenuates the development of B lymphocytes and αβ lineage T lymphocytes in a p53-dependent manner, while sparing other lineages, including γδ T cells ^19,20^. Rpl22 also regulates the emergence of embryonic HSC by controlling the translation of *Smad1* mRNA ^21^. Finally, *RPL22* mutations and deletions have been observed in human T acute lymphoblastic leukemia (T-ALL), and Rpl22 loss promotes lymphoma development in a mouse model of T-ALL, suggesting that it functions as a tumor suppressor ^22^.

The ability of Rpl22 to regulate lymphoid development and transformation, as well as fetal HSC emergence, led us to investigate whether Rpl22 also plays a role in regulating the function of adult HSC and their potential for transformation into myeloid malignancy. Here we show that reduced expression of *RPL22* is seen in human MDS and AML, and, is linked to poor outcomes. This association appears to be causal since Rpl22 loss in mice predisposes to transformation, and does so not though alterations in global protein synthesis, but rather through alterations in HSC metabolism and ultimately through Lin28b-mediated promotion of lipid synthesis in the resulting leukemias.

## RESULTS

### *RPL22* expression is reduced in human MDS and AML

To assess the changes in RP expression levels in human myeloid neoplasms, we performed gene expression analysis of CD34^+^ hematopoietic progenitor cells of 183 MDS patients. This analysis revealed that *RPL22* is one of the most significantly reduced RP in this disease (**Fig. 1A,B**). We also observed that reduced *RPL22* expression was associated with lower hemoglobin (Hb) levels, a marker of MDS severity (**Fig. 1C**)^23^. Since MDS often progresses to AML, we evaluated *RPL22* expression in bulk tumor from AML patients using data obtained from the Beat AML consortium and TCGA^24^. Strikingly, *RPL22* expression was significantly decreased amongst most AML patients compared to healthy controls (**Fig. 1D and S1A)**. We further examined *RPL22* expression levels in the immature cells from which this disease originates, by stringently sorting hematopoietic stem and progenitor cell (HSPC) populations from patients diagnosed with AML (**Fig. 1E-G**)^25^. From these analyses, we observed a significant reduction of *RPL22* mRNA in the leukemia initiating populations comprising long-term HSC (LT-HSC; Lin^-^CD34^+^CD38^-^CD90^+^), short-term HSC (ST-HSC; Lin^-^CD34^+^CD38^-^CD90^-^), and granulocyte-macrophage progenitors (GMP; Lin^-^CD34^+^CD38^+^CD123^+^CD45RA^+^) of patients with higher risk AML (associated with complex karyotype; CK), compared to identically sorted cells from age-matched, healthy controls (**Fig. 1E-G**) ^26,27^. Moreover, reduced expression of *RPL22* was associated with more aggressive AML, since the patients with low *RPL22* expression (bottom quartile) from the TCGA AML cohort exhibited reduced overall survival (**Fig. 1H**). Using fluorescent *in situ* hybridization (FISH) analysis in another independent cohort of MDS/AML samples, we found that the *RPL22* locus was more frequently deleted in progenitor cells from both MDS and AML patients (**Fig. 1I**). Approximately 40% of MDS patients and 27% of AML patients showed evidence of increased *RPL22* deletion, with more of these patients being represented in the high-risk MDS and secondary AML groups **(Fig. 1J)**^28^. Collectively, these data indicate that *RPL22* expression is frequently reduced in MDS/AML patients, including in their HSPC, and that low *RPL22* expression is associated with reduced survival, and thus more aggressive myeloid disease.

**Fig. 1:**
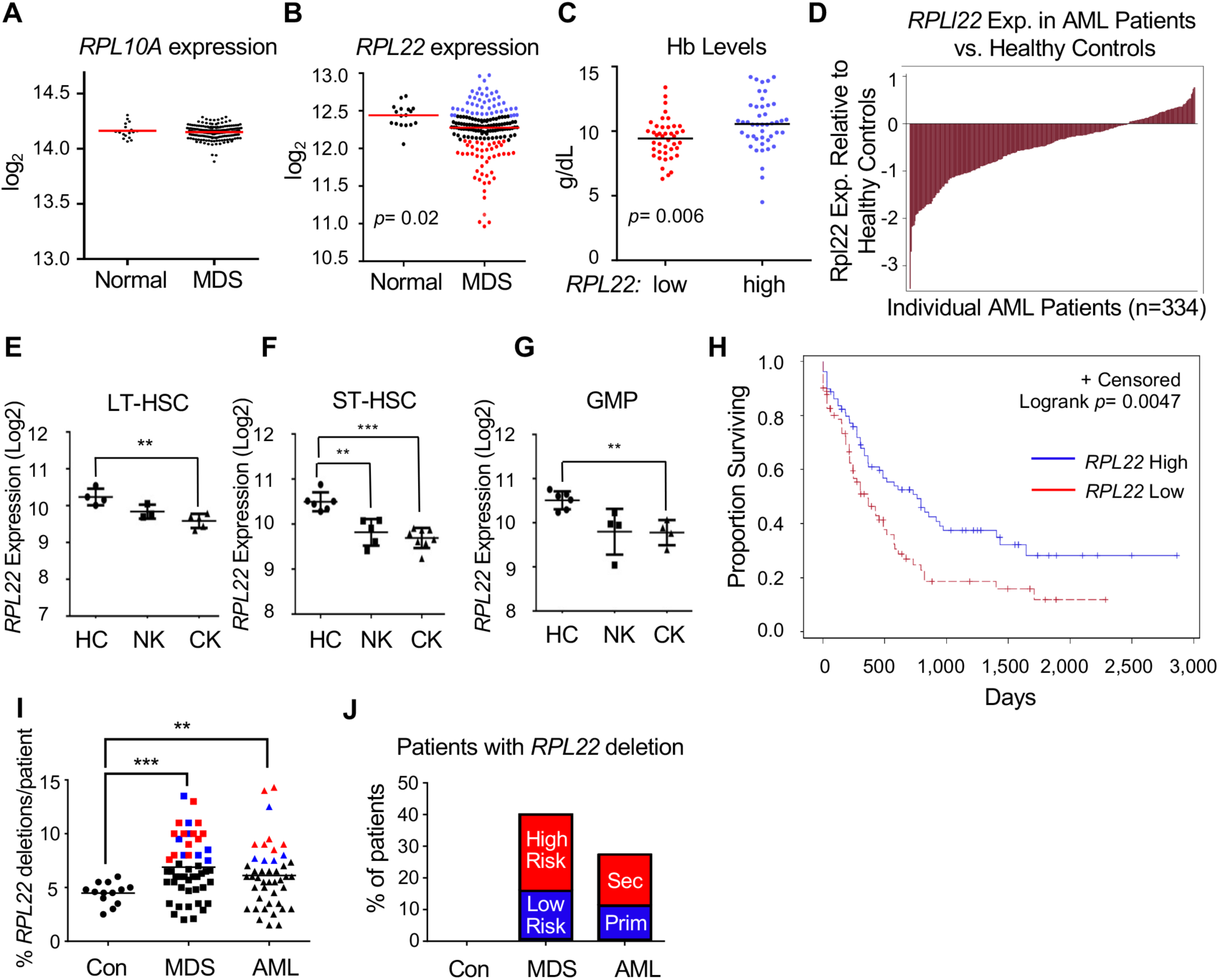
Reduced *RPL22* expression correlates with more severe MDS and AML. **(A-B)** Expression levels of *RPL10A* and *RPL22* mRNA *in* CD34+ cells from MDS patients (*n*=183). **(C)** Hemoglobin (Hb) levels of MDS patients with low (bottom quartile) or high (top quartile) *RPL22* expression. **(D)** Waterfall plot of *RPL22* expression in leukocytes from AML patients and healthy controls enrolled in the Beat AML consortium. **(E-G)** *RPL22* mRNA levels in sorted **(E)** LT-HSC (Lin^-^, CD34+, CD38-, CD90+), **(F)** ST-HSC (Lin-, CD34+, CD38-, CD90-), and **(G)** GMP (Lin-, CD34+, CD38+, CD123+, CD45RA+) from AML bone marrow samples (n=12) compared to age-matched healthy controls (HC; n=4)). Cytogenetic abnormalities are depicted as NK (Normal Karyotype) or CK (Complex Karyotype). **(H)** Graph of survival of AML patients from the TCGA database, with patients with low *RPL22* expression (bottom quartile) compared to those with high expression (top quartile). The survival curve was analyzed for significance using the Mantel-Cox log-rank test. **(I-J)** FISH analysis of deletion of the *RPL22* locus in bone marrow cells from 85 MDS and AML patients. **(I)** Dot-plot representing the fraction of CD34+ cells in which the *RPL22* locus was deleted. Patients in which the frequency of *RPL22* deletion is 3 S.D. greater than the mean of healthy controls are colored, with blue and red colors representing low and high risk MDS, respectively (middle), or primary and secondary AML, respectively (right). **(J)** The distribution of patients with a frequency of *RPL22* deletions greater than 3 S.D. of control were subdivided into low and high risk MDS, and primary and secondary AML, and represented graphically. All graphs were analyzed for significance using the student’s t-test, unless otherwise specified. Error bars represent s.e.m. \*\**p* ≤ 0.01, ***p ≤ 0.001.

### Rpl22-deficient mice display an MDS-like phenotype

To determine whether Rpl22 regulates MDS and AML progression, we assessed whether loss of *Rpl22* disrupted murine hematopoiesis. Similar to MDS patients, *Rpl22*^-/-^ mice displayed a significant reduction in red blood cells (RBC) with increased mean corpuscular volume (MCV), as is observed in macrocytic anemia associated with MDS (**Fig. 2A,B**) ^29^. Rpl22-deficient mice also exhibited evidence of erythroid and myeloid dysplasia (**Fig. S1B,C)**, and an increased frequency of megakaryocytes (**Fig. 2C**). Finally, the bone marrow of Rpl22-deficient mice contained ∼20% more CD34+ cells **(Fig. 2D)** and an expansion of granulocyte-macrophage progenitors (GMP; LK/CD34^+^FCγR^high^; **Fig. 2E**), as is typically observed in higher-risk MDS and AML patients ^30^.

**Fig. 2:**
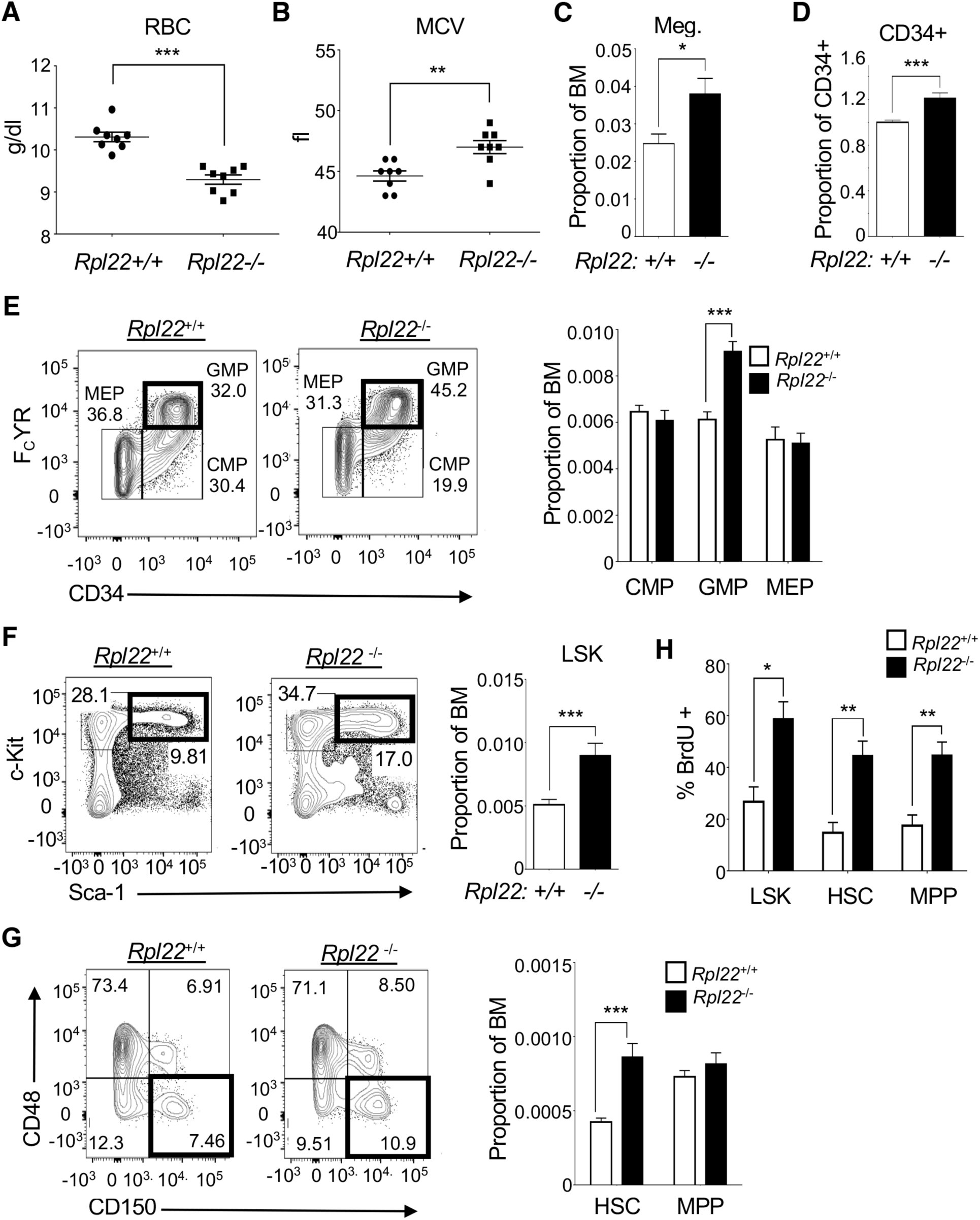
Rpl22-deficient mice exhibit MDS-like characteristics. **(A)** Measurement of red blood cells (RBC; g/dL) in the peripheral blood of *Rpl22*^+/+^ and *Rpl22*^-/-^ mice (*n* = 8). **(B)** Measurement of the mean corpuscular volume (MCV) of RBC from *Rpl22*^+/+^ and *Rpl22*^-/-^mice (*n* = 8). **(C-D)** The proportion of CD41+, FSC-high megakaryocytes (*n* = 9) and CD34+ progenitors (*n* = 18) in the bone marrow of *Rpl22*^+/+^ and *Rpl22*^-/-^ mice was measured by flow cytometry. **(E)** Representative histograms of Lineage- cKit+ (LK) subsets, defined by FcγR and CD34 (including granulocyte-macrophage progenitors, GMP; LK/CD34+/FcγR high) in the bone marrow of *Rpl22*^+/+^ and *Rpl22*^-/-^ mice. Proportions of the indicated populations are represented graphically on the right +/- standard error (*n* = 9). **(F-G)** Representative histograms of LSK and HSC (LSK/CD150+/CD48-) in the bone marrow of *Rpl22*^+/+^ and *Rpl22*^-/-^ mice. The proportions are represented graphically as the mean +/- standard error (*n* = 15). **(H)** Graphical representation of flow cytometric analysis of BrdU incorporation in LSK, HSC, and MPP. (*n* = 4). All graphs were analyzed for significance using the student’s t-test, unless otherwise specified. Error bars represent s.e.m. \**p* ≤ 0.05, \*\**p* ≤ 0.01, \*\*\**p* ≤ 0.001.

Since the alterations in hematopoiesis observed in MDS patients result from impaired function of their expanded HSC population, we sought to determine if these same abnormalities characterized the HSC in *Rpl22*^-/-^ mice ^31,32^. Indeed, both lineage-, Sca-1^+^, c-Kit^+^, (LSK) cells and HSC defined by SLAM markers (LSK/CD48^-^CD150^+^) were increased in the bone marrow of *Rpl22*^-/-^ mice, relative to wild-type littermate controls (**Fig. 2F,G; Fig. S2A,B**) ^33,34^. This expansion was associated with an increased number of proliferating cells, as HSC and other progenitor populations in *Rpl22*^-/-^ mice exhibited greater BrdU incorporation (**Fig. 2H**). *Rpl22-/-* HSC also exhibited a modest increase in Ki-67 staining, compared to *Rpl22*^+/+^ controls (**Fig. S2C;** p=0.06).

Because inactivation of other RP (e.g., *RPS14)* has been reported to impair hematopoiesis by activation of p53, we employed p53-deficient mice to determine if the expansion of pre-malignant HSC in *Rpl22^-/-^* mice was p53-dependent ^13,35^. Interestingly, we found that p53-deficiency failed to suppress the expansion of LSK and HSC observed in Rpl22-deficient mice (**Fig. S2D,E**). Thus, as occurs in MDS patients, Rpl22-deficient mice display an expansion of HSC in the marrow; however, this expansion is not dependent upon p53, as it is not corrected by p53-deficiency. We previously reported that Rpl22 and its paralog Rpl22-Like1 (Rpl22l1 or Like1) antagonistically control the emergence of embryonic HSC, with Rpl22 repressing emergence and Like1 interfering with that repression ^21^. The expansion of adult HSPC in Rpl22-deficient mice is also dependent upon Like1, since ablation of one allele of *Rpl22l1* was sufficient to markedly reduce the number of both LSK and HSC in adult bone marrow **(Fig. S2F-H)**^36,37^.

### Rpl22 loss impairs HSC function

The alterations in hematopoiesis observed in MDS patients have previously been shown to result from dysfunctional HSC ^31,32,38^. To determine if this was also true of the altered hematopoiesis observed in Rpl22-deficient mice, we performed competitive bone marrow transplantation assays using allotype-marked (CD45.2) *Rpl22*^-/-^ or *Rpl22*^+/+^ HSC combined with whole bone marrow competitor (WBM, CD45.1). Analysis of peripheral blood 20 weeks after transplantation revealed that *Rpl22*^-/-^ HSC were significantly less capable of reconstituting hematopoiesis than *Rpl22*^+/+^ HSC, as *Rpl22^-/-^* HSC produced very few peripheral blood leukocytes (**Fig. 3A**). Moreover, the few leukocytes that were produced by Rpl22-deficient HSC were profoundly biased toward the myeloid fate (CD11b^+^Gr-1^+^; **Fig. 3B**), consistent with the basal myeloid-bias observed in the donor *Rpl22*^-/-^ mice and in MDS patients ^39^.

**Fig. 3:**
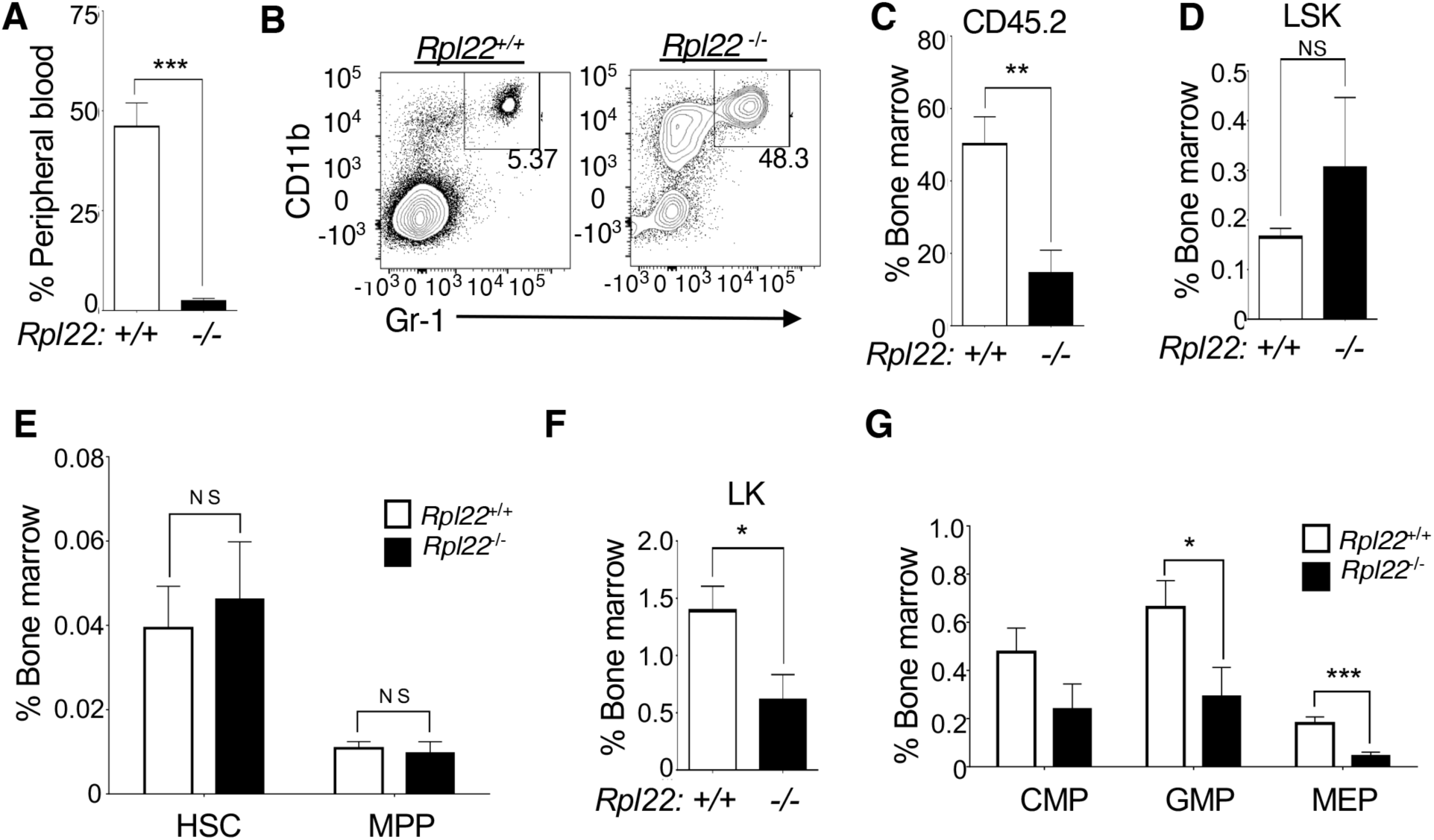
HSC from Rpl22-deficient mice display altered stem cell function. **(A-G)** Competitive transplantation of 300 CD45.2 HSC with 200,000 CD45.1 competitor WBM cells. All data shown represent analysis at 20 weeks post-transplantation (*n* = 10). **(A)** Contribution of CD45.2 *Rpl22*^+/+^ and *Rpl22*^-/-^ HSC to the peripheral blood following competitive transplantation. **(B)** Representative histogram of myeloid lineage cells in the peripheral blood of transplanted mice. **(C-G)** Contribution of CD45.2 *Rpl22*^+/+^ and *Rpl22*^-/-^ HSC to the following bone marrow populations after competitive transplantation: **(C)** total bone marrow cellularity; **(D)** LSK; **(E)** HSC and MPP; **(F)** LK; and **(G)** LK subsets, CMP, GMP, and MEP. All graphs were analyzed for significance using the student’s t-test. Error bars represent standard error of the mean (SEM). \**p* ≤ 0.05, \*\**p* ≤ 0.01, \*\*\**p* ≤ 0.001.

The failure of *Rpl22*^-/-^ HSCs to reconstitute hematopoiesis could have resulted from their failure to engraft, or, alternatively, from failure to produce mature hematopoietic cell lineages following engraftment. To evaluate the extent of engraftment by Rpl22-deficient HSC, we assessed donor chimerism in the bone marrow of transplanted mice. While *Rpl22*^-/-^ HSC contributed less to overall bone marrow cellularity than did *Rpl22^+/+^*HSC (**Fig. 3C**), the contribution of *Rpl22^-/-^* HSC to the recipient bone marrow was nearly 2-times (1.8 fold) greater than to peripheral blood (**Fig. S2I**). Together these data suggest that the MDS-like phenotype observed in *Rpl22*^-/-^ mice results from HSC dysfunction, where Rpl22-deficient HSC are capable of self-renewal, as evidenced by their ability to engraft and expand in the bone marrow, but are impaired in their ability to produce lineage-committed downstream progenitors or peripheral leukocytes.

To identify the stages of hematopoiesis impaired by Rpl22-deficiency, we assessed the contribution of *Rpl22*^-/-^ HSC to hematopoietic progenitor populations, relative to that of *Rpl22^+/+^* HSC. Interestingly, transplanted *Rpl22*^-/-^ HSCs produced LSK, HSC, and multipotent progenitors (MPP, LSK/CD150^-^CD48^-^) as effectively as *Rpl22*^+/+^ HSCs (**Figs. 3D,E and S2J**). However, development of *Rpl22^-/-^* progenitors beyond the MPP stage was more severely impaired, as LK cells, and all of their subsets (GMP; LK/CD34^-^FCγR^low^ megakaryocyte-erythroid progenitors or MEP; and LK/CD34^+^FCγR^low^ common myeloid progenitors or CMP), were reduced relative to mice transplanted with *Rpl22^+/+^* HSC, with GMP and MEP exhibiting the most profound reductions (**Fig. 3F,G**). Consequently, Rpl22-deficient HSC are capable of engraftment, expansion, and generation of MPP, but are impaired in their ability to give rise to downstream, committed progenitors.

### Rpl22-deficiency promotes the development of leukemia

Since self-renewal without the capacity to differentiate characterizes a premalignant state, we reasoned that *Rpl22*^-/-^ HSC should be predisposed to leukemic transformation ^40^. Consequently, we assessed whether *Rpl22*^-/-^ HSC were predisposed to transformation using the MLL-AF9 oncogene knock in model of AML ^41,42^. Indeed, MLL-AF9 knock-in mice exhibited splenomegaly comprising Mac1+ leukemic cells **(Fig. S3A)**, with the *Rpl22^-/-^*mice exhibiting a far greater leukemic burden at 15 weeks of age, as evidenced by splenic weight (**Fig. 4A**). Furthermore, Rpl22-deficient MLL-AF9 knock in mice developed and succumbed to leukemia much more rapidly than their *Rpl22*^+/+^ counterparts (**Fig. 4B**).

**Fig. 4:**
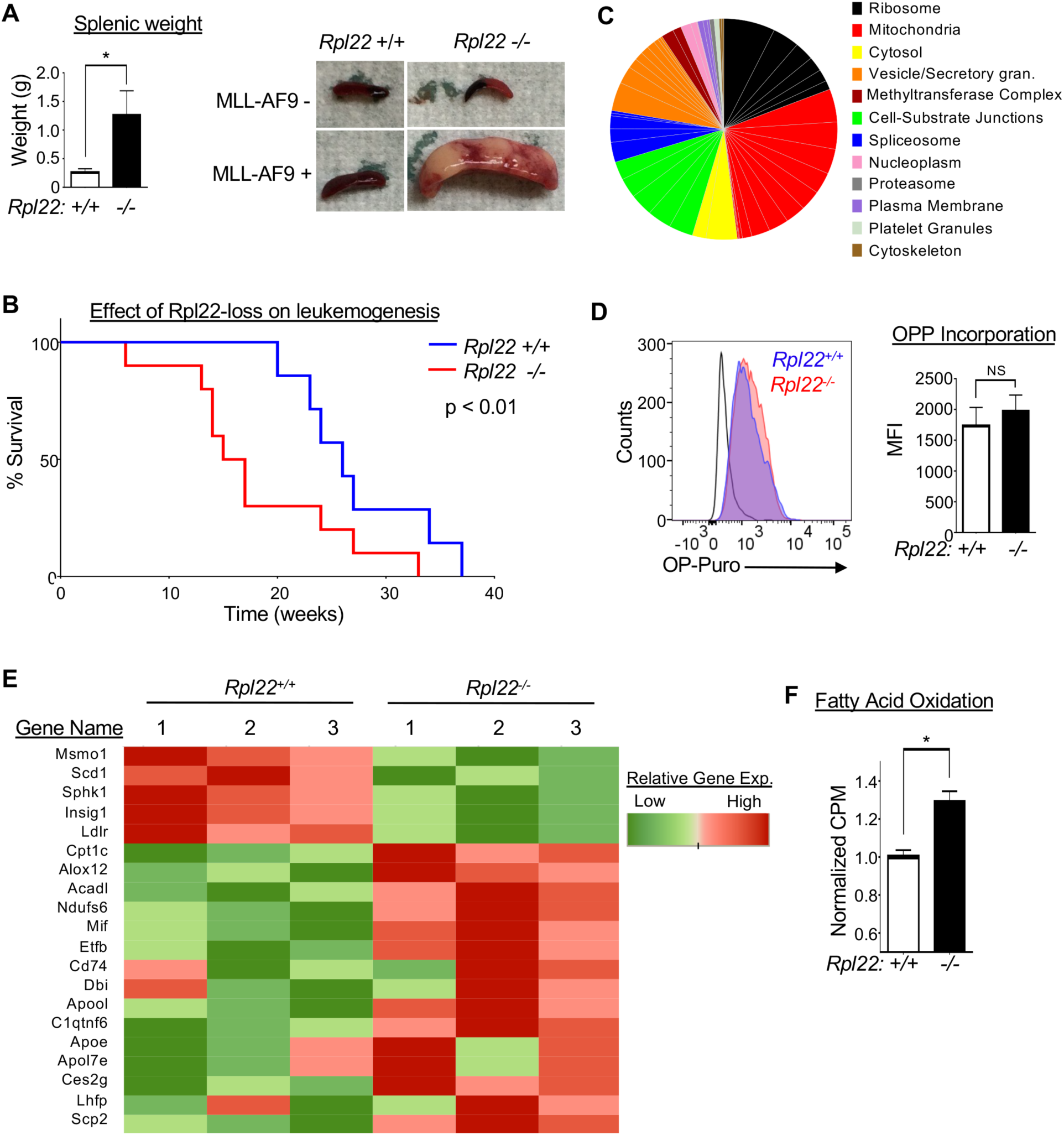
Rpl22 deficiency facilitates leukemogenesis. **(A)** Disease burden on *MLL-AF9* knock in *Rpl22*^+/+^ and *Rpl22*^-/-^ mice was measured at 15 weeks of age (*n* = 5). Representative spleens are depicted along with a graphical representation of the mean +/- SEM of splenic weights. (p < 0.05) **(B)** Survival analysis of *MLL-AF9* knock in *Rpl22*-/- (*n* = 10) and *Rpl22*+/+ (*n* = 7) mice. Significance was determined using a Mantel-Cox test. **(C)** Protein synthesis was measured in HSC by assessing the amount O-propargyl-puromycin (OPP) incorporated into nascent polypeptides by flow cytometry after exposure to OPP for 1h. A representative FACS histogram of OPP incorporation by HSC is depicted and expressed graphically as the mean +/- SEM of the mean fluorescence intensity (MFI). **(D)** Gene ontology (GO) analysis of genes differentially expressed by *Rpl22*^-/-^ HSC. The differentially expressed genes were identified by RNA-Seq and subjected to GO analysis using the GO Cellular Components Database. Significant gene sets were categorized and represented as a pie chart with the size of the wedges reflecting the EnrichR combined score for a given GO category. **(E)** Heat map depicting the expression levels in HSC of genes involved in fatty acid metabolism (n = 3). **(F)** Measurement of fatty acid oxidation (FAO) using C14-labeled palmitate in *Rpl22*^+/+^ and *Rpl22*^-/-^ LSK. FAO activity is depicted graphically as the mean +/- SEM of triplicate measurements. All graphs were analyzed for significance using the student’s t-test. \**p* ≤ 0.05

### Rpl22-deficiency alters the metabolism of HSC and resulting leukemias

To determine how Rpl22-deficiency impairs HSC function and enhances their transformation potential, we performed whole transcriptome analysis on HSC from *Rpl22*^+/+^ and *Rpl22*^-/-^ mice. From this analysis, we found that Rpl22-deficiency altered the expression of more than 500 genes (**Fig. S3B** and **Table S1**). Gene ontology (GO) analysis using the Cellular Components Database revealed that Rpl22-deficiency most significantly affects the expression of clusters of genes associated with the ribosome and mitochondria (**Fig. 4C and Table S2).** Because HSC function has been reported to be impaired by reductions in protein synthesis, and because ribosomal protein mutations are linked to significant decreases in global protein synthesis, we sought to determine if Rpl22-deficiency was impairing HSC function by attenuating protein synthesis ^8,43,44^. To test this possibility, we measured protein synthesis in HSC *in situ*, by monitoring O-propargyl-puromycin (OPP) incorporation into nascent polypeptides using flow cytometry ^43,44^. Surprisingly, we observed that the incorporation of OPP into newly synthesized proteins by *Rpl22*^-/-^ HSC (red) was equivalent to that incorporated by *Rpl22*^+/+^ HSC (blue) (**Fig. 4D**). Therefore, in contrast to what has been observed for HSC upon inactivation of other RP, this analysis revealed the unexpected finding that the alteration of HSC function caused by Rpl22-deficiency is not associated with an attenuation of global protein synthesis ^8,13,43,45^.

Because global protein synthesis was not impaired by Rpl22 loss, we sought to determine if altered mitochondrial function was contributing to the impaired function of Rpl22-deficient HSC, since mitochondrial function, particularly oxidative phosphorylation and fatty acid oxidation (FAO), was another GO class exhibiting an altered expression signature in Rpl22-deficient HSC (**Fig. 4C,E**; **Table S2**). To determine if these changes in gene expression altered mitochondrial function, we performed metabolic analysis on *Rpl22*^+/+^ and *Rpl22*^-/-^ LSK cells. We observed no differences in mitochondrial biomass or in extracellular acidification rate (ECAR), a surrogate measure of glycolysis (**Fig. S4A,B**). Consistent with the lack of change in ECAR, pAMPK, ATP, and lactate levels were also unaffected by Rpl22-deficiency **(Fig. S4C)**. *Rpl22*^-/-^ LSK did exhibit reduced oxygen consumption rate (OCR), suggesting a reduction in cellular capacity to mediate oxidative respiration (**Fig. S4D**). The reduction in oxygen consumption is associated with reduced expression of a key component of the electron transport chain, *Atp5e*, which is required for oxygen consumption by electron transport complex (ETC) IV (**Table S1)**^46^. Interestingly, the expression of several genes involved in fatty acid metabolism was increased in Rpl22-deficient HSC (**Fig. 4E**). These alterations in gene expression are noteworthy because increased dependence on fatty acid oxidation (FAO) has been implicated in both HSC self-renewal, leukemic stem cell function, and cancer stem cell function ^47–49^. Consistent with the increased expression of genes involved in lipid oxidation, FAO was increased in *Rpl22*^-/-^ LSK (**Fig. 4F**). It should be noted that attenuating FAO using the FAO inhibitor, Etomoxir ^50,51^, did not restore oxygen consumption in *Rpl22-/-* HSPC **(Fig. S4D)**.

Since increased FAO has been implicated in the self-renewal of HSC, we asked if explanted Rpl22^-/-^ HSCs exhibited a prolonged retention of their SLAM-marked HSC phenotype (LSK, CD150^+^, CD48^-^) upon culture *in vitro* ^47^. Indeed, whereas a significant proportion of *Rpl22*^+/+^ HSC lost their HSC phenotype after two days in culture, it was fully retained by *Rpl22*^-/-^ HSC **(Fig 5A)**. Moreover, retention of the HSC phenotype by *Rpl22*^-/-^HSC was dependent on FAO, as it was abrogated by inhibition of FAO using Etomoxir (**Fig 5A)** ^50,51^. Blockade of FAO was not cytotoxic to *Rpl22*^-/-^ HSC, but instead induced their differentiation into MPP **(Fig 5B)**. *Rpl22*^-/-^ HSPC also exhibited enhanced function, as they retained a greater capacity to form colonies in methylcellulose following serial passage (**Fig. 5C)**. This capability was also attenuated by pharmacologic inhibition of

**Fig. 5:**
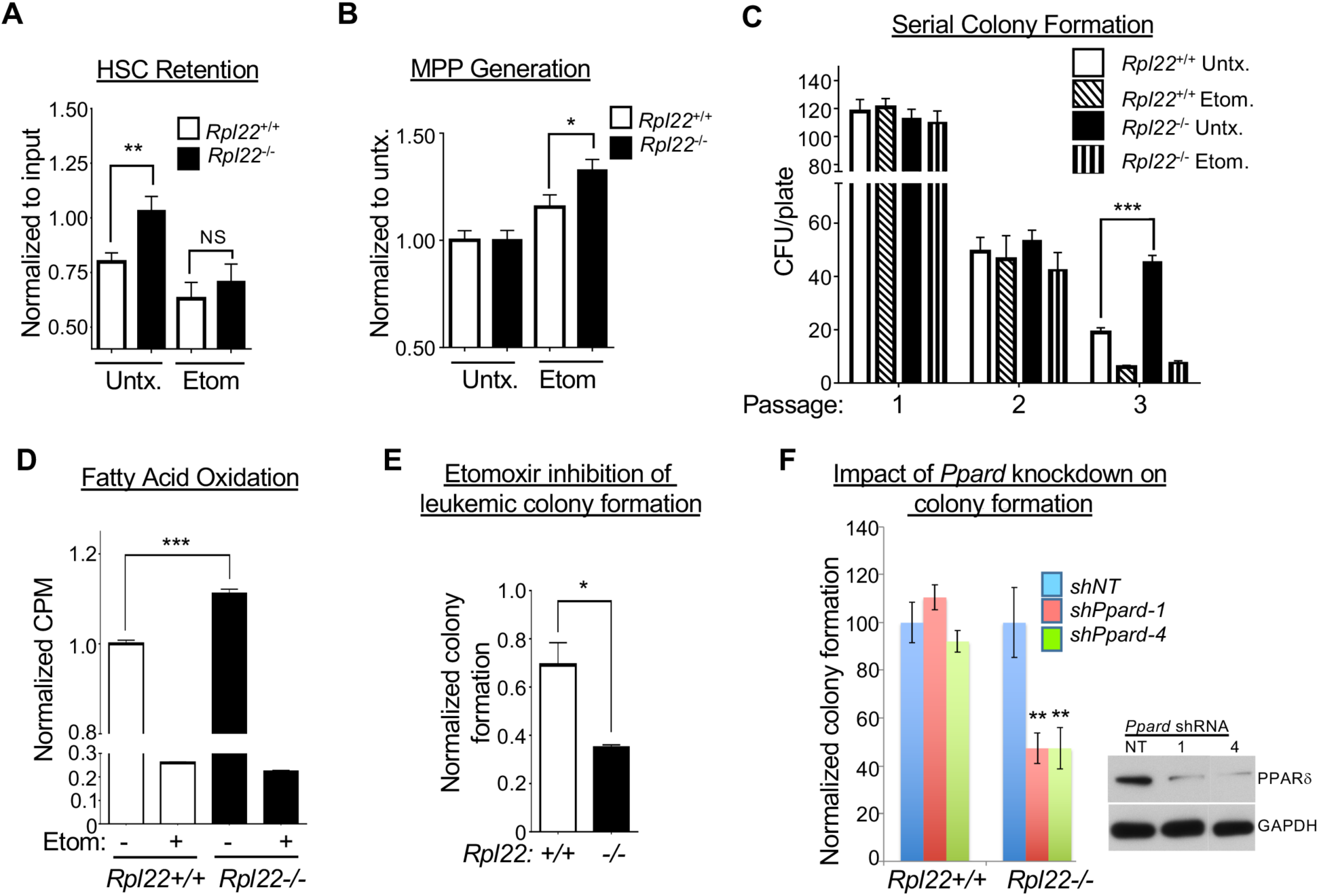
The enhanced retention of progenitor activity by Rpl22-deficient HSPC depends on FAO. **(A-B)** Flow cytometric assessment of the retention of the HSC immunophenotype (CD150+CD48-) or differentiation into MPP following two days of culture in methylcellulose in the presence or absence of 200μM Etomoxir (*n* = 9). **(C)** Serial colony forming ability of *Rpl22*^+/+^ and *Rpl22*^-/-^ HSPCs treated with vehicle (DMSO) or Etomoxir. Mean +/- SEM of colonies of triplicate cultures serially passaged three times is depicted graphically. **(D)** Measurement of FAO using C14-labeled palmitate in MLL-AF9 transformed LSK from *Rpl22*^+/+^ and *Rpl22*^-/-^ mice. Triplicate measurements of FAO activity are depicted graphically as the mean +/- SEM. **(E)** Effect of Etomoxir treatment on colony formation by MLL-AF9 transformed LSK cells from *Rpl22*^+/+^ and *Rpl22*^-/-^ mice. Triplicate measurements of colony formation in methylcellulose are depicted graphically as the mean +/- SEM. **(F)** PPARδ loss selectively attenuates colony formation by *Rpl22-/-* leukemias. MLL-AF9 transformed leukemias from *Rpl22*^+/+^ and *Rpl22*^-/-^ mice were transduced with shRNA targeting *Ppard* following which their colony forming capacity in methylcellulose was assessed. The mean +/- SEM of triplicate measurements is depicted graphically and the effect of the shRNA on PPARδ expression was assessed on B16 melanoma cells by immunoblotting. All graphs were analyzed for significance using the student’s t-test. Error bars represent standard error of the mean (SEM). \**p* ≤ 0.05, \*\**p* ≤ 0.01, ***p≤ 0.001.

FAO (**Figs. 5C and S4E**). Together these findings suggest that the increased self-renewal and inability of *Rpl22*^-/-^ HSC to support the development of downstream committed progeny results from an alteration in cellular metabolism. Specifically, *Rpl22*^-/-^ HSC exhibit enhanced FAO, which promotes self-renewal ^47^.

Since increased FAO was required for the enhanced self-renewal exhibited by *Rpl22*^-/-^ HSC, we next asked if enhanced FAO also contributed to increased leukemic potential of the Rpl22-deficient MLL-AF9 leukemia cells. We found that, similar to non-transformed HSCs, MLL-AF9 transformed LSK from *Rpl22*^-/-^ mice exhibited increased FAO (**Fig. 5D**); The *Rpl22-/-* leukemia did not exhibit the reduction in oxygen consumption observed in primary Rpl22-deficient LSK, suggesting that this defect is not retained during transformation **(Fig. S4F).** The increase in FAO appears to play an important role in maintaining the leukemic potential of *Rpl22-/-* leukemia cells, since treatment with the FAO inhibitor, Etomoxir, was far more effective in inhibiting colony formation by *Rpl22*^-/-^leukemia cells than by their *Rpl22^+/+^* counterparts (**Fig. 5E**). Moreover, the ability of Rpl22-deficient leukemias to form colonies was preferentially-dependent upon the transcription factor PPARδ, a master regulator of FAO that has been implicated in the ability of FAO to promote HSC renewal **(Fig. 5F)**^47^. Indeed, shRNA knockdown of *Ppard* selectively attenuated colony formation by *Rpl22-/-* leukemias **(Fig. 5F)**, indicating that Rpl22 loss promotes leukemic colony formation by upregulating FAO in a PPARδ-dependent manner.

### Rpl22 regulation of FAO and leukemia pathogenesis

Because Rpl22 is an RNA binding protein, we hypothesized that Rpl22 might control FAO in HSC by modulating the activity of mRNA targets encoding regulators of this process. Using M-fold to predict RNA secondary structure, we found that several of the transcripts encoding regulators of FAO bear the consensus stem-loop structure recognized by Rpl22 (**Fig. S5A-E)**. Amongst these differentially expressed targets **(Fig. 4E)**, arachidonate lipoxygenase-12 (Alox12) was of particular interest, both because there were two consensus Rpl22 binding sites in the coding regions of both mouse *Alox12* and human *ALOX12* (**Fig. S5B,F),** and because Alox12 converts polyunsaturated fatty acids to the activating ligands for a master transcriptional regulator of FAO, PPARδ ^52,53^. Analysis of Alox12 protein and mRNA levels revealed that the increase in Alox12 protein levels exceeded that of its mRNA **(Fig. 6A)**, suggesting post-transcriptional regulation. The increase in Alox12 expression was accompanied by increased generation of the arachidonic acid breakdown product it generates, 12S-HETE **(Fig. 6B).** We next wished to determine how the expression of Alox12 was controlled by Rpl22. To determine if Rpl22 bound to the two consensus binding sites in the coding sequence of *Alox12* mRNA, we performed electrophoretic-mobility shift analysis (EMSA) **(Fig. 6C).** EMSA analysis confirmed that Rpl22 bound strongly to positive control RNA, EBER1, but not negative control EBER2 ^54^. Moreover, Rpl22 also bound to the second predicted binding site (BS2) of *Alox12*, but not to the first predicted binding site (BS1) or to NBS, an RNA sequence between BS1 and BS2 that lacks the Rpl22 target motif (**Fig. 6C**). In addition to its ability to bind to *Alox12* mRNA, Rpl22 is capable of post-transcriptionally repressing the expression of Alox12 upon ectopic expression in Rpl22-deficient mouse embryonic fibroblasts (MEF) **(Fig. 6D).** Rpl22 appears to be regulating Alox12 by controlling its inclusion in actively translating polysomes, since Rpl22 reintroduction into Rpl22-deficient MEF displaces *Alox12* mRNA for heavy polysomes **(Fig. 6E).** Moreover, appending the Rpl22 binding site in *Alox12* mRNA (BS2) to a heterologous mRNA (GFP) conferred responsiveness to Rpl22 regulation **(Figs. 6F and S5G),** collectively, indicating that Rpl22 controls HSC function by directly binding *Alox12* mRNA and regulating Alox12 protein expression. To assess whether Alox12 is the critical link between Rpl22-deficiency and elevated FAO, we crossed *Rpl22^-/-^* mice to Alox12-deficiency (*Rpl22^-/-^Alox12^-/-^)*^55^ and assessed FAO in HSPC from these mice using C^14^-palmitate catabolism **(Fig. 6G).** While *Rpl22-/-* HSPC clearly exhibited elevated FAO activity as measured by C^14^-palmitate catabolism, indicating that Rpl22 loss elevates FAO, FAO was not attenuated by the genetic ablation of *Alox12* **(Fig. 6G)**. Together, these data demonstrate that despite Alox12 elevation and the increased presence of its 12S-HETE product, the Rpl22 target Alox12 is not responsible to the enhanced FAO observed in *Rpl22-/-* HSPC.

**Figure 6.**
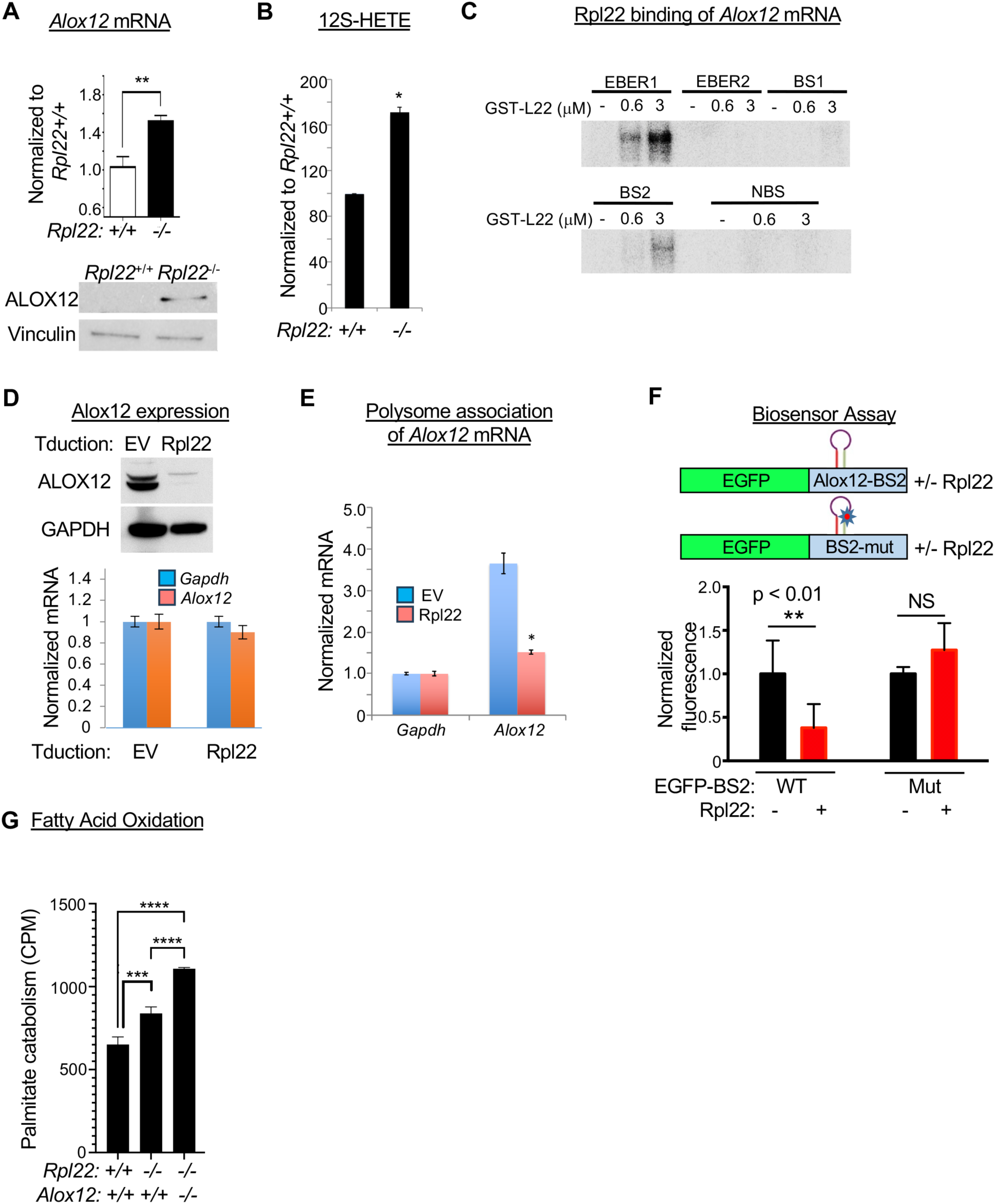
Alox12 is an Rpl22 target but is not responsible for elevated lipid metabolism. **(A)** Q-PCR and Immunoblot analysis of *Alox12* mRNA and protein expression, respectively, in *Rpl22*+/+ and *Rpl22*-/- HSC and LSK. Triplicate qRT-PCR measurements are represented graphically as mean +/- SEM, with the expression level in *Rpl22-/-* HSC normalized to that in *Rpl22+/+* HSC (n=3). Vinculin was used as an immunoblotting loading control. **(B)** The levels of 12S-HETE in *Rpl22+/+* and *Rpl22-/-*LSK were measured in triplicate by ELISA. The mean level +/- SEM was normalized to that in *Rpl22+/+* LSK and depicted graphically. **(C)** Assessment of GST-Rpl22 binding to the indicated fragments of *Alox12* mRNA (BS1, binding site 1; BS2, binding site 2; and NBS, non-binding site) as measured by EMSA analysis. *EBER1* and *EBER2* RNA served as positive and negative controls, respectively. **(D)** Repression of Alox12 expression by Rpl22. *Rpl22-/-* MEF ectopically expressing the coding region of murine *Alox12* were transduced with either the empty vector (EV) control or pMiG-Rpl22, following which the effect on Alox12 expression was assessed by immunoblotting and qRT-PCR as in **(A). (E)** Control of *Alox12* mRNA inclusion in polysomes. Cells from **(D)** were fractionated on a continuous sucrose gradient, which separated cellular RNA into polysome-associated and free fractions. The amount of free and polysome associated *Alox12* and *Gapdh* RNA was quantified by qRT-PCR, and normalized to that in the EV transduced control, as in **(D)**. **(F)** Transfer of Rpl22-responsiveness to mRNA encoding GFP using the Rpl22-binding site in *Alox12* mRNA. *Gfp-*constructs appended in-frame with intact or mutated *Alox12* BS2 were injected into zebrafish embryos at the 1 cell stage alone or with mRNA encoding Rpl22 **(Fig. S5G)**. After 10h, GFP fluorescence was quantified and depicted graphically after normalization to the fluorescence intensity of the intact BS2 construct in control injected fish. All data are representative of 3 experiments performed, unless otherwise stated. Significance for pairwise tests was assessed using the student’s t-test while multiple comparisons were assessed by one-way ANOVA. Error bars represent standard error of the mean (SEM). \**p* ≤ 0.05, \*\**p* ≤ 0.01, \*\*\**p*≤ 0.001, \*\*\*\**p*≤ 0.0001 **(G)** Measurement of FAO using C14-labeled palmitate on LSK from mice of the indicated genotypes. Triplicate measurements of FAO activity are depicted graphically as the mean +/- SD. *** p<0.0004; **** p<0.0001;

### Role of Lin28b in promoting the pathogenesis of Rpl22-/- leukemias

To gain insight into the molecular basis for the control of lipid metabolism by Rpl22 and to make a broader assessment of the regulation of leukemogenesis by Rpl22, we performed RNA-Seq on *Rpl22+/+* and *Rpl22-/-* leukemias that developed in the MLL-AF9 transgenic mice **(Figs. 7A and S6A).** The analysis revealed 2671 differentially expressed genes, with 1219 being induced and 1452 being repressed in *Rpl22-/-* leukemias **(Fig. 7A; Table S3).** Genes upregulated in *Rpl22-/-* leukemias include many that have been previously implicated in AML pathogenesis **(Fig. 7A).** *Prtn3* and *Laptm4b* are reported to promote leukemogenesis by regulating STAT3 signaling, while *Dock1* is a guanine nucleotide exchange factor thought to promote leukemogenesis through Notch activation^56–59^. Key transcriptional regulators are also induced in *Rpl22-/-* leukemias including *Mansc1* (MN1), which serves as a critical transcriptional co-factor in MLL rearranged leukemias and *Six1*, which promotes leukemia by regulating the expression of glycolytic genes **(Fig. 7A)**^60,61^. Also among the induced genes are critical metabolic regulators of leukemia progression **(Table S3)**. *Cyb561* promotes leukemogenesis through ROS induction and lncRNA *Spehd* regulates oxidative phosphorylation and mitochondrial membrane potential required for stem cell function^62,63^. Many of the induced genes have also been identified as essential for leukemia survival in CRISPR screens **(Figs. 7B).** Finally, the repressed genes in *Rpl22-/-* leukemias are similarly implicated in leukemia pathogenesis including *Asxl2, Cdkn2a,* and *Hpse2,* which have been found to be frequently mutated in leukemias **(Figs. 7A and S6B)**^64–67^. Among the diverse differentially regulated genes in *Rpl22-/-* leukemias, 10% are linked to metabolism, with fatty acid metabolism being the most enriched, upregulated metabolic pathway **(Figs. 7C,D and S6C-E).** Together, these data indicate that Rpl22 regulates leukemic progression by broadly regulating the expression of critical regulators falling in numerous gene ontology groups, but particularly in lipid metabolism.

**Fig. 7:**
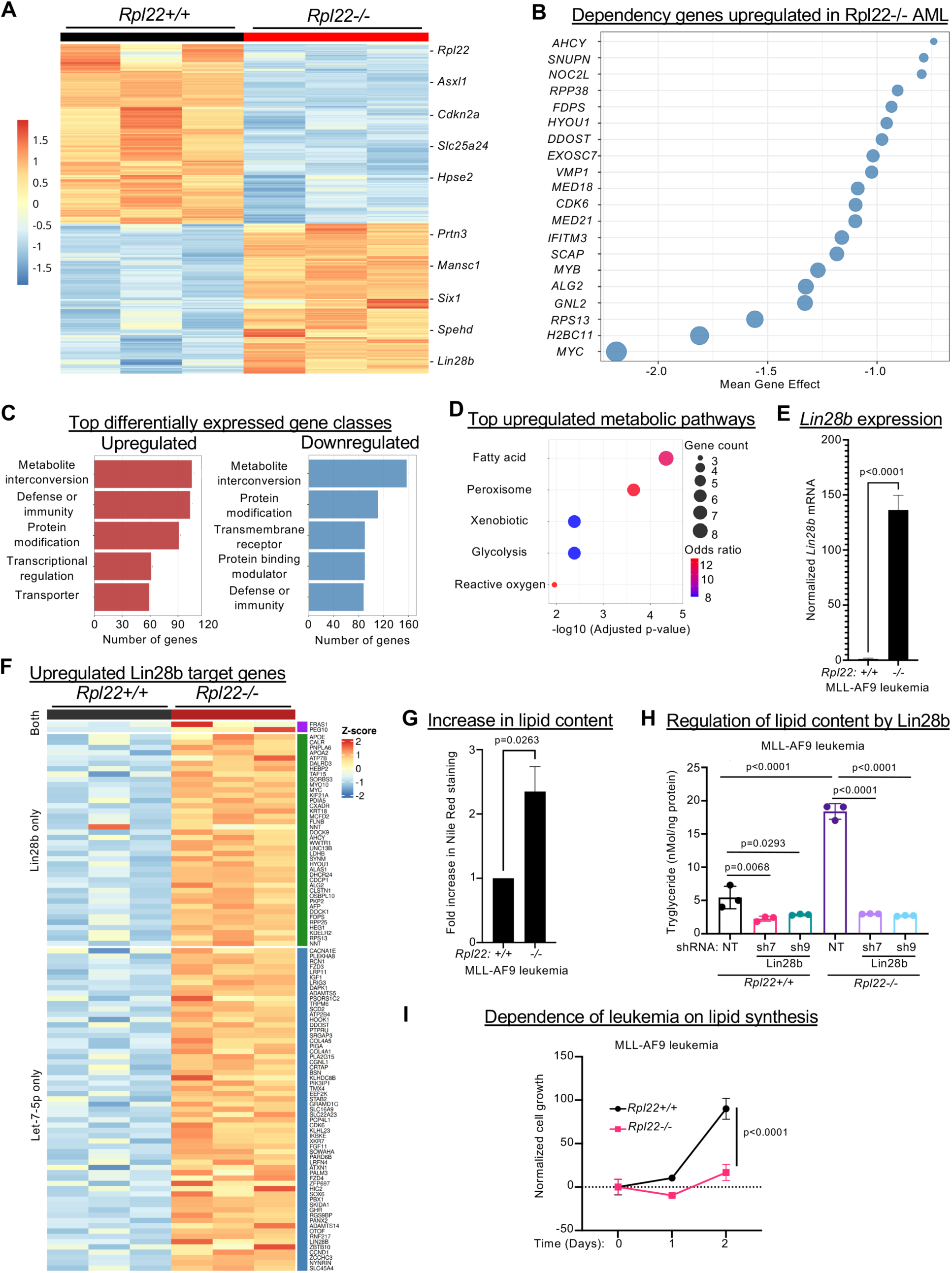
Rpl22-deficient leukemia cells are dependent on elevated lipid synthesis. **(A)** Leukemia explants from MLL-AF9 knockin *Rpl22^+/+^* (M82) and *Rpl22^-/-^* (M109) mice were analyzed by RNA-Seq. A heat map of 2,671 differentially expressed genes is displayed. **(B)** Top 20 upregulated essential genes in *Rpl22^-/-^* leukemias, identified using the CRISPR DEPMAP CERES dataset across human leukemia cell lines and displayed as a bubble plot of the impact of their genetic disruption on leukemia survival. **(C)** Differentially expressed genes were subjected to pathway analysis (PANTHER classification) with the top upregulated and downregulated pathways displayed. The number of differentially expressed genes per pathway is indicated. **(D)** Bubble plot of the top five Reactome pathways depicting the most upregulated metabolic pathways in *Rpl22^-/-^* MLL-AF9 leukemias. **(E)** Quantitative PCR analysis of *Lin28b* mRNA expression in *Rpl22+/+* and *Rpl22-/-* MLL-AF9 leukemias. Statistical significance was calculated using a Two-tailed T test with Welch’s Correction. **(F)** Heat map of 103 Lin28b gene targets induced in *Rpl22^-/-^* MLL-AF9 leukemias, subdivided based on whether they are direct Lin28b targets or are indirectly regulated through Lin28b modulation of Let7 micro-RNAs (mIR). **(G)** Lipid content of *Rpl22^+/+^* and *Rpl22^-/-^* leukemias as measured by Nile Red staining. Statistical significance was calculated using a Two-tailed t test with Welch’s Correction. **(H)** Triacylglycerol (TG) levels were measured on detergent extracts of *Rpl22^+/+^* and *Rpl22^-/-^*leukemias using a colorimetric assay quantifying oxidized glycerol liberated by lipase digestion. Mean +/- SD of triplicate measures of nMol TG per ng protein were expressed graphically. Statistical significance was determined using One Way ANOVA. **(I)** The effect of inhibiting TG synthesis using DGAT1 inhibitor (DGAT1-IN-1) at 25μM on the growth of *Rpl22^+/+^* and *Rpl22^-/-^* leukemias was assessed by MTT assay. Triplicate measures were expressed graphically as mean +/- SD for each different drug concentration. Statistical significance was determined by two way ANOVA.

Lin28b has been implicated as a key regulator of both leukemia pathogenesis and lipid metabolism, raising the possibility that Lin28b induction may play a key role in regulating lipid metabolism in *Rpl22-/-* leukemias^68–75^. We have previously shown that Rpl22 loss promotes development of T acute lymphoblastic leukemia (T-ALL) through Lin28b induction^22^. *Lin28b* is also one of the most upregulated genes in leukemias that arose in the MLL-AF9 Tg *Rpl22-/-* mice **(Figs. 7A,E).** Importantly, of the 1219 genes upregulated in Rpl22-deficient leukemias, nearly 10% (102) are either direct Lin28b targets or indirect targets regulated through Lin28b effects on Let-7 micro-RNAs (mIR) **(Fig. 7F and Table S4)**^76,77^. These targets include many that are required for leukemia survival, such as Myc, Rps13, and CDK6 **(Fig. S7A)** as well as those implicated in lipid metabolism, **(Fig. S7B)**. Along with the expression signature, Nile Red staining revealed that *Rpl22-/-* leukemias displayed greater lipid content **(Fig. 7G).** To determine if the increased lipid content resulted from Lin28b induction, we knocked Lin28b down using shRNA **(Fig. S7C).** Indeed, Lin28b knockdown reduced triacylglycerol (TG) synthesis, preferentially in *Rpl22-/-* leukemias **(Fig. 7H).** Finally, inhibition of Acyl-CoA:diacylglycerol acyltransferase 1 (DAGAT1)^78^, a critical enzyme in TG synthesis^79^, resulted in preferential attenuation of the survival of *Rpl22-/-* leukemias **(Fig. 7I)**, indicating that Lin28b-mediated promotion of lipid synthesis is critical for pathogenesis of *Rpl22-/-* leukemias.

## DISCUSSION

In this report, we provide evidence that RP-insufficiency alters HSC function and increases the predisposition to leukemia through a novel mechanism that does not involve attenuation of global protein synthesis. Instead, HSC function is altered because of the dysregulated expression of selected Rpl22 gene targets that impact metabolism. *RPL22* expression is reduced in human MDS and AML, including at the stem cell level, and is associated with reduced survival. The link between reduced *RPL22* expression and poor survival is also observed in a mouse leukemia model lacking Rpl22. As indicated, the mechanistic link between Rpl22 loss and enhanced leukemogenic potential is distinct from that of other cases of RP insufficiency in that it does not involve attenuation of global protein synthesis ^8,13,43^. Instead, *Rpl22* inactivation creates a premalignant state characterized by enhanced HSC self-renewal and impaired generation of downstream committed progenitors, by altering metabolism. Specifically, Rpl22-loss decreases oxygen consumption and increases the dependence of HSC and leukemias on FAO. These observations are consistent with previous reports demonstrating that enhanced FAO promotes HSC self-renewal and cancer stem cell activity, including leukemia stem cells ^47–49,80,81^. The increased FAO is associated with induction of a regulator of FAO that is a direct Rpl22 target, Alox12^52,82^, but Alox12 is not responsible for increased FAO since its loss does not return FAO to baseline. Instead, we identified induction of the stemness factor Lin28b in the resulting leukemias that is a key driver of the enhanced lipid synthesis (TG) that supports Rpl22-deficient leukemia survival. Together, these observations indicate that Rpl22 controls HSC function and transformation potential through effects on metabolism.

Mutations in RP have previously been reported to perturb hematopoiesis and predispose to transformation^2,3,13,83–89^; however, Rpl22-deficiency appears to regulate hematopoiesis and predispose to transformation in a manner that is fundamentally different from that of other RP mutations, since Rpl22 is not essential for life and Rpl22-deficiency does not result in detectable alterations in ribosome function^19^. Nevertheless, Rpl22-deficiency does disrupt normal hematopoiesis at multiple stages, including the perturbation of HSC function reported here^19–22,37,90–93^. Two modes of action have been proposed to explain how insufficiency of an RNA-binding RP might regulate hematopoiesis: through effects on the ribosome itself or through extraribosomal activity^9,11,94–96^. This remains a controversial issue, as there is evidence in support of both modes of action. Effects on the ribosome itself can impact HSC function by altering the overall protein synthesis rate, since normal HSC function is critically-dependent on maintaining the rate of protein synthesis within a very narrow range^43,97,98^. Rpl22-deficiency does not reduce the global protein synthesis rate, suggesting that Rpl22-deficiency does not disrupt HSC function by changing global protein synthesis. Another way RP-insufficiency could affect the ribosome is that RP insufficiency may produce ribosomes with altered protein composition (specialized ribosomes), which exhibit distinct capacities to translate mRNA species bearing specific primary sequence motifs or secondary structural features ^3,12,99,100^. While evidence supporting this perspective has been reported^92^, an opposing model suggests that RP insufficiency attenuates the translation of a selected class of mRNA species by decreasing the number of available ribosomes, rather than by altering their composition^8^. The completely distinct mode by which RP can regulate processes is by leaving the ribosome and functioning in a physically separated manner, referred to as “extra-ribosomal” function^96^. Clear support for this model has also been reported, including the well documented ability of Rpl13a to dissociate from the ribosome following interferon signaling and to bind to selected mRNA targets, thereby regulating their translation^101^. Our analysis does not definitively distinguish the mode of action through which Rpl22 functions; however, our evidence aligns best with the extra-ribosomal model. Indeed, Rpl22 assembles into the ribosome as a monomer with a single RNA binding face^102,103^, which is bound to the 28S rRNA, precluding it from simultaneously having direct interaction with another RNA target. Consequently, direct interaction of Rpl22 with mRNA targets, such as *Alox12,* is only possible when the RNA-binding helices of Rpl22 are free from the 28S rRNA, as is the case for the Rpl22 pool that is physically separated from the ribosome.

We have identified a number of targets through which Rpl22 regulates development, including hematopoiesis^21,37^. We previously reported that the antagonistic balance between Rpl22 and its highly homologous paralog, Rpl22-Like1 (Rpl22l1 or Like1), controls the emergence of embryonic HSC by directly binding and controlling the translation of *Smad1* mRNA^21^. Rpl22 regulation of the behavior of adult stem cells is also dependent upon Like1, since Like1-insufficiency attenuates the HSC expansion observed in Rpl22-deficient mice, though this presumably does not involve effects on Smad1 expression, since Smad1 is dispensable in adult HSC ^104^. We determined that the Rpl22-Like1 balance plays a critical role in controlling gastrulation by binding and regulating the splicing of many pre-mRNA targets, including Smad2, an essential molecular effector of gastrulation^37^. Rpl22 regulates traversal of pre-receptor checkpoint stages for B and T lymphocytes ^20,92^. At least in T cells, the regulation of this transition is mediated through control of the unfolded protein response (UPR) or endoplasmic reticulum (ER) stress signaling^92^. The lineage restricted requirement for Rpl22 in regulating ER stress signaling appears to be limited to cells experiencing unusually abrupt transitions from quiescence to rapid proliferation, as is observed at the pre-receptor checkpoints ^105^. This does not appear to be relevant in cells that undergo less abrupt proliferative transitions, such as γδ T cell progenitors or adult HSC^19^. We report here that dysregulation of FAO by Rpl22 loss results in HSC dysfunction and is required for the survival of Rpl22-deficient leukemias, which is impaired by knockdown of the master regulator of FAO, PPARδ^47^. Rpl22 directly binds and regulates the translation of mRNA encoding Alox12, which has been implicated in FAO^52,53^; however, Alox12 is not responsible for the increased FAO observed in Rpl22-deficient HSPC and so we do not think Alox12 plays a role in the predisposition to transformation displayed by Rpl22-deficient HSPC. Instead, the upregulation of FAO may result from dysregulation of any number of other genes, including Electron Transfer Flavoprotein-β (ETF-β), which both regulates FAO and supports AML pathogenesis^106,107^.

These and other observations underscore the critical role that alterations in lipid metabolism play in solid and hematologic malignancies, as well as in the control of HSC function ^47–49,80,108^. Nevertheless, the molecular basis by which enhanced FAO supports the function of normal HSC and leukemia cells generally, and specifically in the context of Rpl22 loss, remains to be established. There are three potential non-mutually, exclusive processes that might contribute to these altered behaviors. First, FAO is capable of generating energy through the contribution of NADH and FADH to the mitochondrial electron transport chain (ETC) to generate ATP ^109^. However, *Rpl22-/-* HSPC exhibited a reduction in oxygen consumption rate, a surrogate measure or aerobic respiration, suggesting that the generation of energy was unlikely to be responsible. The reduction in oxygen consumption is likely to result from reduced expression of *ATP5e*, which is important for the consumption of oxygen by ETC complex IV ^46^. This is consistent with our observation that *Atp5e* expression is no longer reduced in the *Rpl22-/-* leukemias, which exhibit OCR levels equivalent to their *Rpl22+/+* counterparts (data not shown). Second, FAO might enhance the survival of *Rpl22-/-* HSPC, since FAO has been reported to modulate the function of Bcl2 family members ^110^. This could explain how FAO promotes the survival of *Rpl22-/-* leukemias. Finally, FAO also generates substantial quantities of acetyl-CoA, which can enter the Krebs cycle to generate citrate and contribute to NADPH-producing reactions. Alternatively, the acetyl-CoA can be released to the cytosol where it could alter cell behavior through the acetylation of both cytosolic and nuclear proteins ^111,112^. Indeed, our preliminary proteomic analysis has revealed that Rpl22-deficient leukemias exhibit profound alterations in protein acetylation (data not shown). Efforts are in progress to distinguish among these possibilities.

Our data suggest that upregulation of Lin28b, which plays a broader role in lipid metabolism, supports the pathogenesis of Rpl22-deficient MLL-AF9 transgenic AML through induction of TG synthesis. Lin28b is primarily expressed in fetal progenitors where it serves as a master regulator of the fetal hematopoietic program, since its ectopic expression in adult progenitors is sufficient to recapitulate many aspects of fetal hematopoiesis^70,113^. A key question is how Rpl22-deficiency results in Lin28b induction. While we have shown that Rpl22 can acutely regulate Lin28b expression^22^, bioinformatic assessment of the *Lin28b* gene did not identify any Rpl22-binding motifs in either the exonic or intronic sequences, strongly suggesting *Lin28*b is not a direct Rpl22 target (data not shown). We have previously shown that Lin28b upregulation is critical for the pathogenesis of Rpl22- thymic lymphomas driven by transgenic expression of myristoylated-Akt2 and in this case Lin28b induction was dependent upon NFκB^22^. NFκB does not appear to be responsible for Lin28b induction in the Rpl22-deficient myeloid leukemia cells since the RNA-seq analysis did not identify an NFκB signature. Myc is both a Lin28b target and a regulator of its expression^71,114–116^ and so it remains unclear whether the increased expression of Myc in the Rpl22-deficient AML is the cause or a result of Lin28b induction. Interestingly, the expression of Lin28b is not elevated an adult Rpl22-deficient HSC. This is perhaps not surprising because Lin28b is primarily expressed in fetal progenitors^117^, but does raise the possibility that the progenitor population for the resulting Rpl22-deficient MLL-AF9 transgenic leukemias may be of fetal origin. Irrespective of the gestational origin of the leukemia initiating cell, Rpl22-deficiency cooperates with MLL-AF9 to lead to both Lin28b induction and the disabling of the recently reported tumor suppressive activity of Lin28b in postnatal MLL-based leukemogenesis^69^. Lin28b has been implicated in the pathogenesis of a variety of cancers, including

AML bearing MLL translocations^68,71,74,118^. Lin28b is an RNA-binding protein that regulates processes, including carcinogenesis, either by direct binding to mRNA targets or indirectly by regulating the processing of *Let7* family micro-RNAs (mIR)^118–120^. Rpl22-deficient leukemias exhibit increased expression of a large number of direct Lin28b targets as well as those regulated through *Let7* action, including Myc, which has been implicated in the Lin28b-mediated pathogenesis of numerous cancer types^116,121,122^. Lin28b is also a well-known regulator of cellular metabolism^70,72,75^ and has been reported to support cancer progression by supporting *de novo* fatty acid synthesis^73^. Lin28b does so through direct binding to and control of the translation of mRNAs encoding critical regulators of the synthesis of multiple TG species, SREBP-1 and SCAP^73^. TG content is significant greater in *Rpl22-/-* leukemias and they are more dependent on TG synthesis for survival than their *Rpl22+/+* counterparts. While we did not observe induced mRNA encoding SREBP-1 in *Rpl22-/-* leukemias, this is not surprising since their regulation by Lin28b is posttranscriptional. However, SCAP and SCD2, both targets of SREBP-1 and critical regulators of TG synthesis, were induced in *Rpl22-/-* leukemias^123,124^, suggesting that SREBP-1 activation is also responsible for the augmented production of TG in Rpl22-deficient leukemias. SCAP was also identified as an Rpl22-regulated gene upon which leukemia survival depends **(Fig. 7B).**

Taken together, our observations indicate that the RP, Rpl22, employs a novel mode of action to regulate the transformation potential of HSC that does not involve altering global protein synthesis, but is instead focused on the control of cellular lipid metabolism. Importantly, perturbations in lipid metabolism are increasingly understood to represent therapeutic vulnerabilities in AML ^49,125–130^; however, past efforts exploit this therapeutically in patients have been limited by liver toxicity ^131,132^. Our findings suggest that targeting TG synthesis may represent a more fruitful approach^133–135^, particularly in those patients with *RPL22*-insufficiency.

## ACKNOWLEDGEMENTS

This work was supported by NIH grants R37AI110985, P30CA006927, an appropriation from the Commonwealth of Pennsylvania, the Leukemia and Lymphoma Society, and the Bishop Fund. We are grateful to the Wiest laboratory for helpful suggestions and Dr. Avinash Bhandoola, for critical evaluation of the manuscript. These studies were supported by the following core facilities at Fox Chase: Cell Culture, Flow Cytometry, Biostatistics and Bioinformatics, Genomics, and Laboratory Animal.

## AUTHOR CONTRIBUTIONS

B.H., D.K.S. designed and performed most of the experiments, interpreted data and wrote the manuscript. S.P.F., S.R.S, B.T., and S.P. analyzed the RNA-Seq data. M.V., J.P., R.K., O.G., J.B., A.P., K.R., C.M., K.P., M.W., J.W., J.T., M.H., B.K., Y.Z., M.R., R.P.,S.S., B.A., S.A. and U.S. helped with experiments. In collaboration with A.V, and S.S., D.L.W. oversaw the design, execution, and interpretation of the data, and assisted in writing the manuscript.

## STAR METHODS

### KEY RESOURCE TABLE

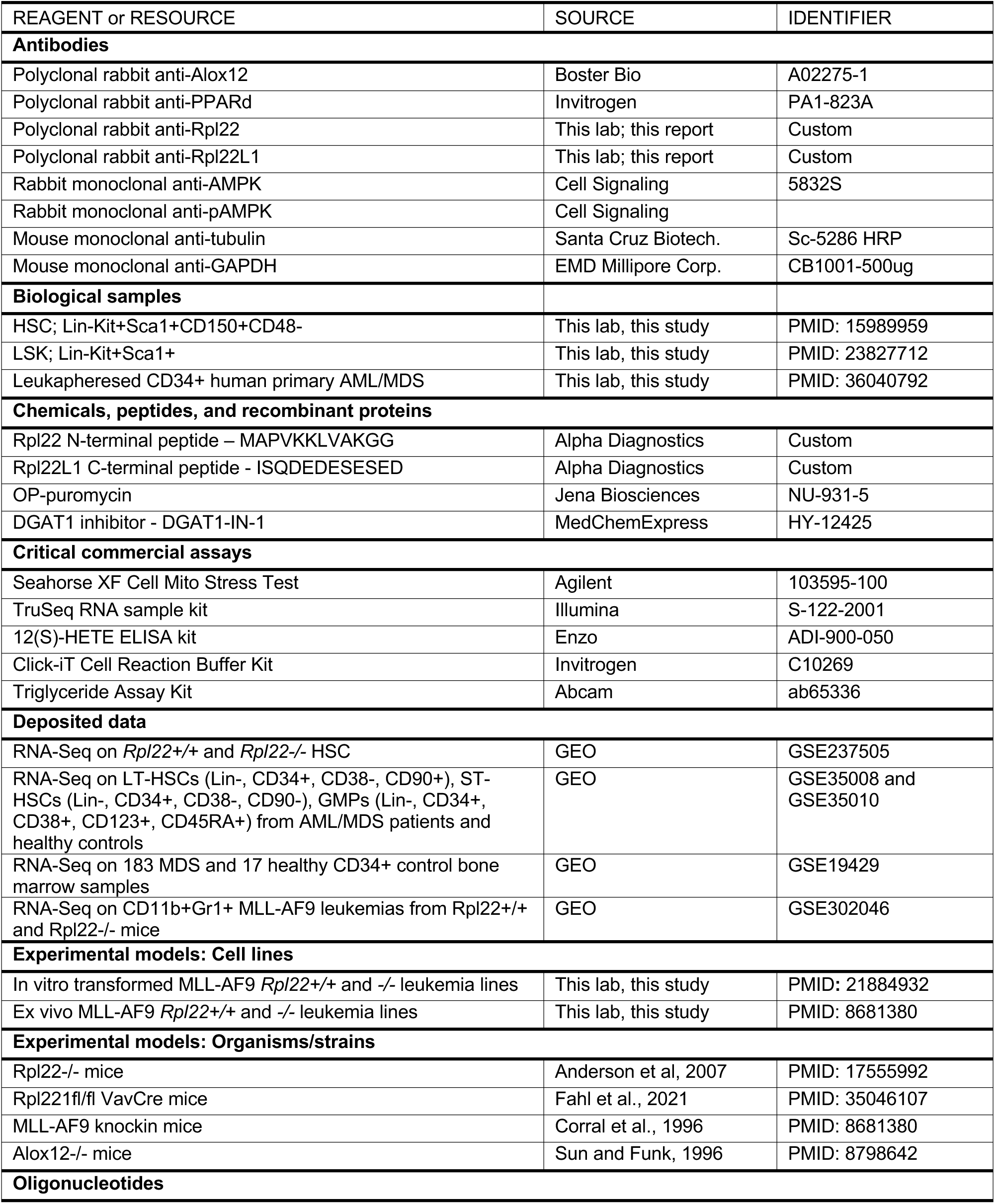
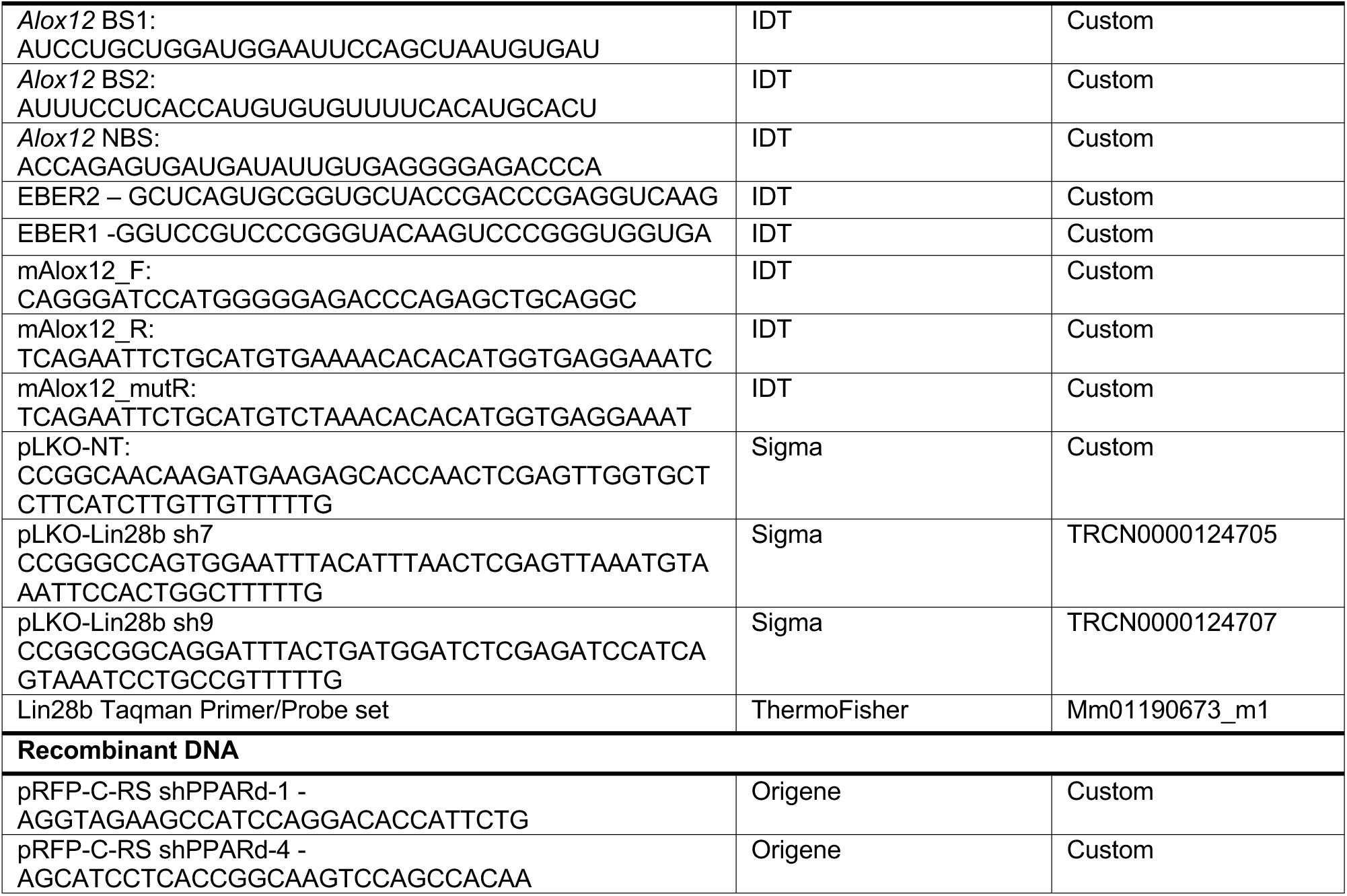

### RESOURCE AVAILABILITY

#### Lead Contact

Further information and requests for resources and reagents should be directed to and will be fulfilled by the Lead Contact, David Wiest

### Materials Availability

All reagents and novel mouse strains used in this study are available upon request from the Lead Contact, David Wiest.

### Data and Code Availability

The RNA-Seq data generated in this study for *Rpl22+/+* and *Rpl22-/-* HSC have been deposited in GEO and can be accessed using GSE237505. Gene expression data on sorted LT-HSCs (Lin-, CD34+, CD38-, CD90+), ST-HSCs (Lin-, CD34+, CD38-, CD90-), GMPs (Lin-, CD34+, CD38+, CD123+, CD45RA+) from AML/MDS patients and healthy controls is deposited in the GEO database (GSE35008 and GSE35010). The RNA-Seq data generated from *Rpl22+/+* and *Rpl22-/-* CD11b+Gr1+ MLL-AF9 leukemias were deposited in the Gene Expression Omnibus database (GSE302046).

## EXPERIMENTAL MODEL AND SUBJECT DETAILS

### Mice

Mice were maintained in the Association for Assessment and Accreditation of Laboratory Animal Care-accredited Laboratory Animal Facility at Fox Chase Cancer Center and were handled in compliance with guidelines established by the Institutional Animal Care and Use Committees. Young adult *Rpl22+/+, Rpl22-/-,* and *Alox12-/-* mice (8-12wks of age) were used in all experiments and had been backcrossed to the C57BL/6 background^19,55^. CD45.1 allotype marked C57BL/6 mice were purchased from Jackson Labs (Bar Harbor, ME). *MLL-AF9* knockin mice were also purchased from Jackson Labs ^42^, backcrossed with Rpl22-deficient mice in our colony, following which survival analysis was performed on littermates. For survival analysis, *MLL-AF9* knockin mice were sacrificed upon developing symptoms of disease (i.e. difficulty breathing, hunched posture, poor grooming, or obvious splenic protuberance) or at specified times for disease burden analysis.

### Patient database and survival data

The expression of ribosomal protein genes and their correlations with clinical parameters were derived from gene expression studies of 183 MDS and 17 healthy CD34+ control bone marrow samples (GSE19429). AML survival curve and Rpl22 expression data was obtained from the Beat AML consortium ^24^. Patient samples used in generation of the survival curve were obtained within 60 days of diagnosis. AML survival data was confirmed with 200 AML samples from TCGA database. Survival curves were calculated by Kaplan Meir analysis. Gene expression data on sorted LT-HSCs (Lin-, CD34+, CD38-, CD90+), ST-HSCs (Lin-, CD34+, CD38-, CD90-), GMPs (Lin-, CD34+, CD38+, CD123+, CD45RA+) from AML/MDS patients and healthy controls is deposited in the GEO database (GSE35008 and GSE35010).

### FISH analysis of the RPL22 locus

A FISH probe for *1p36.2* with the RP11-MI719 BAC was produced commercially by Empire Genomics by nick translation. The TelVysion orange 1q probe was used as a control. Control values for the percent deletion of *RPL22* were established with pooled XY control bone marrow samples (pool of 20 bone marrows which tested negative by New York State and College of American Pathologists (CAP) Guidelines) as well as individual patient samples (patients with anemia and initial lymphoma bone marrow samples – all with normal cytogenetics). The probe was tested in 112 patient samples including patients with low risk MDS, high risk MDS, as well as primary and secondary AML. Over 2600 interphase cells in total were counted for the XY Control samples. At least 200 interphase cells were counted for each of the patient control samples as well as the MDS and AML samples. Metaphase cells were examined when identified. PRISM (http://www.graphpad.com/scientific-software/prism/) software was used to analyze the data for correlations with disease severity and other chromosomal abnormalities.

### Complete blood counts

Peripheral blood was collected into EDTA coated tubes by cardiac puncture. Blood was analyzed using the Abaxis VetScan Hematology Analyzer (Union City, CA) according to the manufacturer’s instructions.

### Immunofluorescence, histology, and immunoblotting

Sternums were formaldehyde fixed, decalcified, and subjected to H&E staining using a standard weak acid protocol. Tissues were paraffin embedded for sectioning. Immunofluorescence analysis was performed on bone marrow sections, bone marrow touch preps, or cytospun cell suspensions. Images were captured using a Nikon E800 upright microscope with BioRad Radiance 2000 confocal scanhead. Primary antibodies were conjugated using AlexaFluor 594 or 488 (Life Technologies) and DAPI was used for nuclear staining. Rabbit polyclonal anti-Rpl22 and anti-Rpl22L1 antibodies were produced by conjugating the N-terminal 12 amino acids of Rpl22 (MAPVKKLVAKGG) and the C-terminal 12 amino acids of Rpl22L1 (ISQDEDESESED) to KLH and immunizing rabbits^136^. Harvested anti-serum was assessed for specificity using samples from *Rpl22-/-* and *Rpl22l1-/-* mice. Immunoblotting was performed as described^21^.

### Flow cytometric analysis and cell sorting of hematopoietic stem and progenitors

Bone Marrow was isolated either by flushing or crushing bones with mortar and pestle. Phenotypic analysis was performed using femurs only. The suspension was passed through a 100 μm filter, subjected to ACK Lysis (pH 7.4) of RBC, and then washed with PBS (without Ca2+ or Mg2+) containing 2% FBS. Enrichment for stem and progenitors was performed by negative selection of mature cells using rat antibodies to mature lineage markers (Gr-1, CD11b, B220, Ter119, and CD3) and goat anti-rat magnetic beads (Qiagen). Cells were stained using standard approaches with the indicated antibodies. PI or DAPI was used to exclude dead cells or for nuclear staining in cell cycle analysis. In assays requiring CD34 staining, cells were stained on ice for a minimum of one hour to ensure appropriate antigen binding. Cells were analyzed for flow cytometry using a BD LSR II and sorted using BD FACSAria II.

### In vivo proliferation analysis of the hematopoietic compartment

Proliferation was assessed by BrdU incorporation. Briefly, mice were injected with BrdU at a dose of 0.15mg/g body weight. Mice were also maintained on water supplemented with 1mg/mL BrdU and 2% sucrose. After 24 hours, mice were sacrificed and bone marrow was isolated. Cells were surfaced stained then fixed and permeabilized for nuclear staining using the Foxp3 Transcription Factor Staining Buffer Set (eBioscience). Cells were DNAse treated at 37°C for 1h and then incubated with anti-BrdU antibody (BioLegend). Cell cycle activity was also confirmed by Ki-67 Staining. Briefly, cells were lineage reduced then stained with a fixable live-dead dye. Lineage reduced cells were then surface stained for HSC markers, fixed, permeabilized using the Foxp3 Transcription Factor Staining Buffer Set, and then stained with anti-Ki-67, (BioLegend), following which they were analyzed by flow cytometry.

### Competitive transplantation

CD45.1 mice were lethally irradiated with a total dose of 11 Gy, split in two doses of 6.5Gy and 4.5Gy, separated by three hours. After 24 hours, mice were injected retro-orbitally with 300 CD45.2+ LSK/CD48-/CD150+ HSC and 200,000 CD45.1+ competitor bone marrow cells delivered in 200μl Hank’s Balanced Salt Solution (HBSS). Engraftment was monitored by retro-orbital bleeding every 4 weeks. Mice were sacrificed after 20 weeks of engraftment and bone marrow was analyzed by flow cytometry.

### In vivo analysis of protein synthesis

Protein synthesis by HSC in vivo was measured with OP-Puro, as previously described ^43,44^. Mice were injected with 50mg/kg OP-Puro and after one hour the mice were sacrificed and bone marrow was isolated. After lineage reduction, cells were stained with GhostDye710, a fixable live/dead dye (Tonbo Biosciences), and analyzed by flow cytometry using the indicated antibodies. After staining, the cells were fixed and permeabilized using a Fix/Perm kit (BD Biosciences), following which the OP-Puro was conjugated to azide-linked AlexaFluor 488 (Life Technologies) using the Click-It Cell Reaction kit (Life Technologies). Alexa 488 fluorescence was then measured by flow cytometry.

### RNA-Seq analysis

HSC were sorted directly into 500μL TriReagent (Sigma-Aldrich), as previously described, following which RNA was isolated according to the manufacturer’s protocol. RNA-Seq libraries were prepared using TruSeq RNA sample kit according to the manufacturer. The RNA-Seq gene set (Table S1) lists mRNA targets differential expressed between *Rpl22*^-/-^ and *Rpl22^+/+^* HSC. RNA-Seq reads were processed for quality issues using FastQC (S.Andrews,http://www.bioinformatics.babraham.ac.uk/projects/fastqc/). Processed reads were aligned to the mouse genome (mm10) using TopHat2 (^137^, following which gene counts were quantified using HTSeq ^138^. The resulting gene counts were used as input for differential expression analysis between Rpl22+/+ and Rpl22-/- HSC using DESeq2 ^139^). The application of a p-value filter of 0.001 resulted in identification of 649 differentially expressed genes. The set of differentially expressed genes was then evaluated using the Mouse Genome Atlas, KEGG, and OMIM Disease Databases through the Enrichr Platform (http://amp.pharm.mssm.edu/Enrichr/.) To compare the genes differentially expressed in HSC upon Rpl22 loss to those modulated by MLL-AF9 expression, we compared our transcriptome data to that published by Stavropoulou *et al.* ^140^, focusing on the 72 hr time point (GSE65384). The expression data was RMA normalized and Limma was used to identify differentially expressed genes ^141,142^. All calculations were done using packages from Bioconductor in the R programming environment ^143^. Heatmaps were generated using the Pheatmap package available through Bioconductor package repository. The RNA-Seq study was deposited in the Gene Expression Omnibus database (GSE237505).

For RNA-Seq on MLL-AF9 transgenic leukemias, explanted CD11b+Gr1+ splenocytes from tumor bearing *Rpl22+/+* and *Rpl22-/-* MLL-AF9 transgenic mice were purified by flow cytometry and processed for RNA isolation using the NucleoZOL reagent (Macherey-Nagel), following the manufacturer’s protocol. RNA (250 ng) underwent poly(A) enrichment, first- and second-strand cDNA synthesis, and library amplification with the Illumina mRNA Stranded Library Kit, per the manufacturer’s instructions. Libraries were pooled, quantified, and loaded at 750 pM onto a NextSeq 2000 flow cell. Sequencing was performed with a 59-10-10-59 (R1-I1-I2-R2) cycle configuration, targeting 30–50 million paired-end reads per sample. Raw reads were trimmed with Trimmomatic to remove Illumina adaptors, and transcript quantification was conducted using Salmon^144^. Downstream analysis—including principal component analysis and differential expression testing—was performed in R using DESeq2. Pathway and gene set enrichment analyses were carried out with EnrichR and GSEA. Plots were generated with ggplot2. The RNA-Seq data were deposited in the Gene Expression Omnibus database (GSE302046). CRISPR dependency scores were obtained from the DepMap portal^145^. Lin28b targets were defined based on CLIP-seq data^76^, and predicted *let-7-5p* targets were retrieved from TargetScan Mouse 8.0^77^.

### Metabolic profiling

100,000 LSK cells were incubated overnight in the presence of C14-palmitate (1.7 μCi), TPO (100 ng/mL), and SCF (100 ng/mL), following which the cells and supernatant were acid precipitated to remove and non-oxidized palmitate. Soluble, oxidized lipid was quantified by liquid scintillation counting. FAO was also assessed on 150,000 MLL-AF9 transformed LSK labeled for 4h. Cellular bioenergetics of LSK cells sorted from Lin- depleted bone marrow cells was determined using the extracellular flux analyzer (XF^96^ analyzer, Seahorse Bioscience). Briefly, Seahorse cell culture analysis plates were coated with CellTak (BD Biosciences) one day before the experiment. LSK and MLL-AF9 transformed LSK cells were suspended in sterile serum-free assay buffer (RPMI1640 supplemented with 5.5 mM D-glucose, 4 mM L-glutamine, and 1 mM pyruvate, pH 7.4) and centrifuged for 5 mins at 1200 rpm to allow them to settle and adhere on the plates. Media was carefully aspirated without disturbing the cells. Fresh assay media was added to control cells. Cells were treated with Etomoxir (100 uM) for 30 minutes at 37 °C and then analyzed by the extracellular flux analyzer according to the manufacturer’s instruction. FCCP, Oligomycin, Rotenone/Actinomycin were injected in the wells following standard protocol of the Agilent Seahorse XF Cell Mito Stress Test.

### Rpl22 regulation of ALOX12 expression

Rpl22 binding sites in mRNA encoding regulators of FAO were identified as described using M-fold software with previously identified stem-loop binding sequence structures^37^. To evaluate predicted binding sites in *Alox12* mRNA, EMSA analysis was performed. 35 nucleotide RNA oligos encompassing binding sites and controls were 5’-end labeled with γ-32P-ATP. 15pmol of each RNA oligo was incubated with 40μCi of γ-32P-ATP and 10U T4 polynucleotide kinase (NEB, M0201S) in 1x T4 polynucleotide kinase buffer (NEB) for 40 min at 37°C. Free ATP was removed using a NucAway spin column (ThermoFisher, AM10070). For EMSA reactions, 5nM of radioactive labeled RNA was added to binding buffer (37.5 mM HEPES (pH 7.9), 75 mM NaCl, 5 mM MgCl2, 0.1mg/ml BSA, 8% glycerol, and 1 μg E.coli tRNA) containing GST-Rpl22 fusion proteins. RNA-binding was assessed by electrophoresis on a non-denaturing gel. Oligo sequences used were as follows:

*Alox12* BS1: AUCCUGCUGGAUGGAAUUCCAGCUAAUGUGAU *Alox12* BS2: AUUUCCUCACCAUGUGUGUUUUCACAUGCACU *Alox12* NBS: ACCAGAGUGAUGAUAUUGUGAGGGGAGACCCA EBER2: GCUCAGUGCGGUGCUACCGACCCGAGGUCAAG EBER1: GGUCCGUCCCGGGUACAAGUCCCGGGUGGUGA

To evaluate the capacity of Rpl22 to regulate ALOX12 expression, the *Alox12* coding region (pLVX-*Alox12-mCherry)* was retrovirally transduced into the *Rpl22-/-* MEF line (KOML3), following which the cells were transduced with either empty vector (pMiG) or *Rpl22* (pMiG-*Rpl22).* Subsequently, the capacity of Rpl22 to regulate ALOX12 protein and mRNA was assessed by immunoblotting and qRT-PCR, respectively, on doubly transduced (mCherry/GFP double positive) cells. To determine the basis by which Rpl22 regulates ALOX12 expression, the aforementioned cells were treated with cycloheximide, following which detergent extracts were subjected to sedimentation on a linear sucrose gradient to separate free mRNA from that being translated in heavy polysomes, as described ^21^. The *Alox12* mRNA content in the free and polysome-associated mRNA pools was then quantified by qRT-PCR and that in Rpl22-expressing cells was normalized to control transduced cells, and the distribution of *Gapdh* mRNA. The capacity of Rpl22 binding sites to transfer Rpl22-responsiveness to a heterologous mRNA target was assessed by appending target sequences with a start codon and fusing them in frame to GFP to generate a biosensor, as described ^18,92^. The constructs were then cloned into pCS2 and transfected into KOML3 cells, following which the GFP+ cells were transduced with empty vector (pMiCherry) or Rpl22 (pMiCherry-*Rpl22).* The effect on GFP levels were assessed by quantifying the mean fluorescence intensity (MFI) using FlowJo software, and mRNA levels were quantified by qRT-PCR in isolated, double-transduced cells. The target sequences employed comprise 141bp fragment of mouse wild *Alox12* containing the Rpl22 binding hairpin loop (BS1-WT) fused in frame with GFP and subcloned into pCS2+. The following primer sets were used:

mAlox12_F: CAGGGATCCATGGGGGAGACCCAGAGCTGCAGGC; mAlox12_R: TCAGAATTCTGCATGTGAAAACACACATGGTGAGGAAATC.

The mutant fragment was generated such that the hairpin loop is not formed using the following reverse primer:

mAlox12_mutR: TCAGAATTCTGCATGTCTAAACACACATGGTGAGGAAAT.

### Generation of MLL-AF9 leukemia lines

LSK were sorted from Rpl22+/+ and Rpl22-/-mice, transduced with MLL-AF9-retroviruses, and serially-passaged through M3434 methylcellulose until stabilized (StemCell Technologies). Cells were then maintained in IMDM supplemented with 10% FBS, 10 ng/mL SCF, 6 ng/mL IL-3, and 5 ng/mL IL-6.

### Colony Formation Assays

For colony formation assays, 500 primary or MLL-AF9 transformed LSK were plated in M3434 Methylcellulose (Stem Cell Technologies) in 35 mm^2^ dishes. Colonies were counted every seven days. For serial replating assays, after seven days cells were isolated by dissolution in media and 10,000 cells were re-passaged. To ectopically express ALOX12, the murine *Alox12* coding sequence was subcloned into the pLVX-IRES-mCherry lentiviral vector. 48h after transduction into *Rpl22+/+* LSK, the mCherry+ cells were purified by cell sorting and subjected to colony formation analysis as above. *Ppard* knockdown in MLL-AF9 leukemias was performed using shRNA in pRFP-C-RS that were obtained from Origene:

shPPARd-1: AGGTAGAAGCCATCCAGGACACCATTCTG shPPARd-4: AGCATCCTCACCGGCAAGTCCAGCCACAA

RFP+ MLL-AF9 leukemias transduced with control (NT) or *Ppard-*targeting shRNA were sorted and then plated to assess the effect on colony formation as described above. For drug treatment with etomoxir, 500 primary LSK or MLL-AF9 transformed LSK cells were resuspended in M3434 Methylcellulose supplemented with 200 μM etomoxir. Cells were plated on 35 mm^2^ plates and colony formation was assessed after seven days. Etomoxir was purchased from Sigma-Aldrich and dissolved as specified by the manufacturer. The drug was stored in the dark at -20C, and thawed only once prior to use. For drug treatment with baicalein, 500 MLL-AF9 transformed LSKs were resuspended in M3434 methylcellulose supplemented with indicated concentrations of drug. Baicalein was purchased from Sigma-Aldrich and fresh dilutions were made prior to each experiment in DMSO. The drug was stored in the dark at -20C.

### Metabolite quantitation

Levels of 12(S)-HETE were measured in triplicate in flow cytometrically sorted LSK from Rpl22^+/+^ and Rpl22^-/-^ mice. The cells were assayed for 12(S)-HETE levels using the 12(S)-HETE ELISA kit from Enzo Life Sciences, Inc (Ann Arbor, MI) according to the manufacturer’s instructions. 25,000 cells per well were loaded to assess 12(S)-HETE production and were normalized to 12(S)-HETE levels in Rpl22^+/+^ LSK. Lactate and ATP were measured using Colorimetric Assay Kits from Biovision (Cat#K607 and Cat#K354, respectively) according to manufacturer’s recommendations. For lactate measurements, LSK cells were lysed in hypotonic lysis buffer (10mM Tris-Cl (pH7.2), 1mM EDTA, 150 mM NaCl, Protease inhibitor, 0.05% Triton X-100), and then the clarified supernatant was used to quantify lactate at 570nm using a standard curve. For ATP measurement, purified LSK and lysed in ATP Assay Buffer, deproteinized using a 10kDa spin column, and used to quantify ATP using a standard curve.

### Lin28b regulation of lipid content and survival

Explanted Rpl22+/+ and Rpl22-/-leukemias from MLL-AF9 transgenic mice were cultured in vitro in 10ng/ml murine stem cell factor, 6ng/ml interleukin-6, and 5ng/ml interleukin-3. Lipid content of these cells was measured by staining with Nile Red. 5 x10^6^ cells were stained in 1ml HBSS containing 3μM Nile Red at 37°C for 10 min, washed with cold HBSS containing 1%BSA, and assessed by flow cytometry using Helix Blue as a viability dye. In addition, triglycerides (TG) were measured in detergent extracts of Rpl22+/+ and Rpl22-/- MLL-AF9 Tg leukemias using the Abcam TG Assay Kit according to manufacturer’s recommendations. The impact of Lin28b knockdown on lipid content and growth of leukemias was assessed by knocking down Lin28b using shRNA.

pLKO (Puromycin resistance cassette replaced with GFP):

NT – CCGGCAACAAGATGAAGAGCACCAACTCGAGTTGGTGCTCTTCATCTTGTTGTTTTTG

sh7 - CCGGGCCAGTGGAATTTACATTTAACTCGAGTTAAATGTAAATTCCACTGGCTTTTTG

sh9 - CCGGCGGCAGGATTTACTGATGGATCTCGAGATCCATCAGTAAATCCTGCCGTTTTTG

Leukemia lines were transduced with the shRNA encoding lentiviral constructs and 48h later GFP-expressing transduced cells were isolated by flow cytometry. Isolated cells were used for the analysis above. The impact of Lin28b shRNA on *Lin28b* expression was assessed by quantitative PCR using a Taqman probe set (Mm01190673_m1; ThermoFisher). The dependence of *Rpl22+/+* and *-/-* leukemias on TG was determined by performing MTT assays according to the manufacturer’s recommendations following treatment with a DGAT1 inhibitor (DGAT1-IN-1).

### Statistical analysis

A two-tailed student’s t-test was used to compare significance between groups in all graphed data. ROUT method was used to identify and confirm clear outliers. Outliers determined by this method are denoted using an asterisk on graphs. Survival-Curves were analyzed using the Mantel-Cox log-rank test. All analyses were performed using Microsoft Excel or GraphPad Prism Software. P-values, Z-scores, or combined scores associated with gene set enrichment analyses were obtained using the built-in function used by the Enrichr Analysis.

