## Supplementary material for "Ribosomal protein control of hematopoietic stem cell transformation through regulation of metabolism": Harris et al., 2025 Supplementary Data .pdf

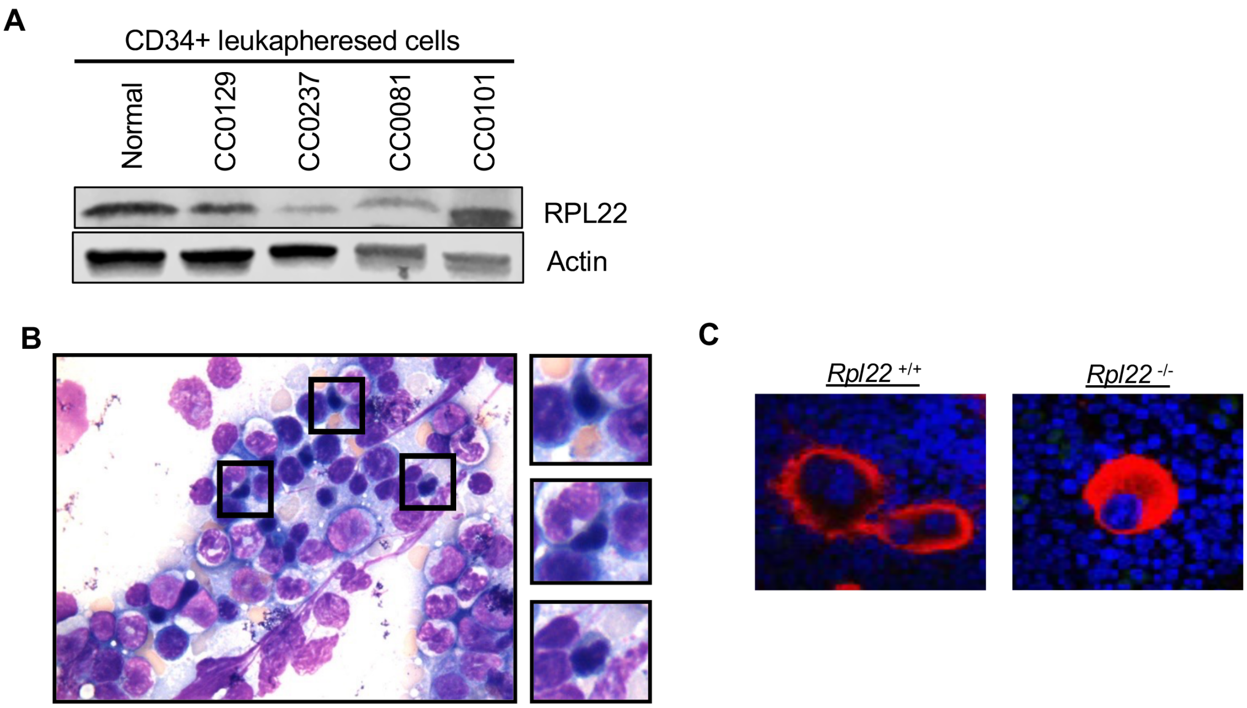


**Figure S1, related to Figure 2. RPL22 insufficiency is associated with aggressive leukemia in humans and dysplasia in mice.**
**(A)** Rpl22 protein expression in leukapheresed cell samples from normal and AML patients. **(B)** H&E staining of *Rpl22*-/- bone marrow. Insets represent images of dysplastic erythroblasts. **(C)** Immunofluorescent staining of bone marrow for megakaryocytes with CD41 (red) and nuclei stained with DAPI (blue).

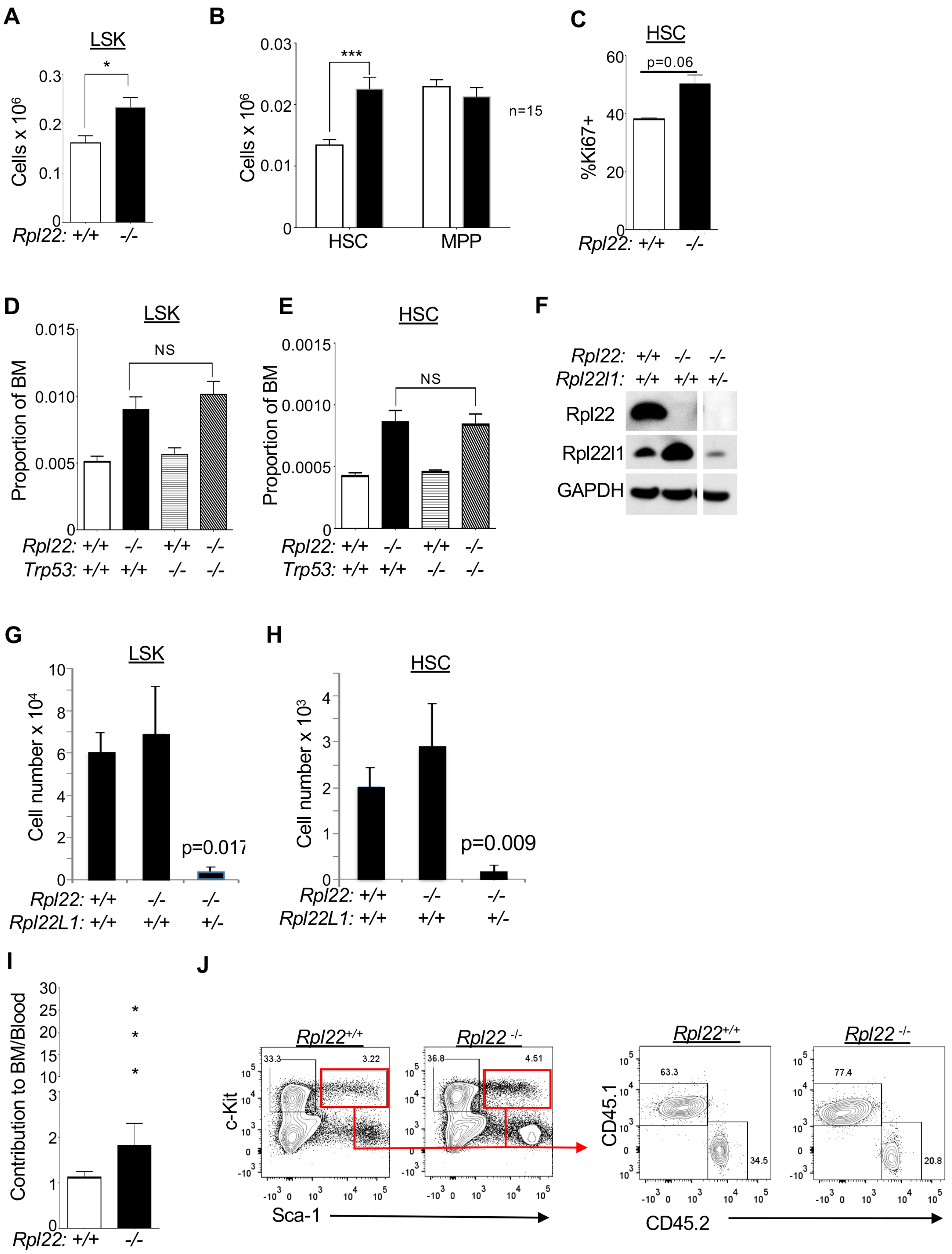


**Figure S2, related to Figure 2: Rpl22 regulation of HSPC is p53-independent. (A-B)** Graphical representation of flow cytometric analysis of numbers of LSK **(A),** HSC, and MPP **(B)**

in the femoral bone marrow of *Rpl22*$^{+/+}$ and *Rpl22*$^{-/-}$ mice (*n* = 15). **(C)** Assessment of cell cycle status of *Rpl22*$^{+/+}$ and *Rpl22*$^{-/-}$ HSC by internal staining for Ki-67. The mean +/- SEM of Ki67+ cells from a representative experiment repeated twice is represented graphically (*n* = 4). **(D-E)** Flow cytometric analysis of the proportion of LSK **(D)** and HSC **(E)** in the bone marrow of *Rpl22*$^{+/+}$*Trp53*$^{+/+}$, *Rpl22*$^{-/-}$*Trp53*$^{+/+}$, *Rpl22*$^{+/+}$*Trp53*$^{-/-}$, and *Rpl22*$^{-/-}$*Trp53*$^{-/-}$ mice (*n* ≥ 6). **(F-H)** *Rpl22l1* was conditionally and monallelically ablated in the bone marrow of Rpl22-/- mice using pIpC-mediated (Mx1-Cre) ablation of the *Rpl22l1*$^{f/f}$ allele. The effect on Like1 protein levels was assessed by immunoblotting of Like1, Rpl22, and GAPDH as loading control **(F).** The effect of *Rpl22l1* ablation on numbers of LSK and HSC **(G,H)** was determined by flow cytometry and depicted graphically as absolute number per mouse (upper and lower leg bones)**.** The mean +/- SEM of representative experiments is represented graphically. All graphs were analyzed for significance using the student's t-test. *$p \leq 0.05$; ***$p \leq 0.001$. **(I)** Ratio of contribution of transplanted CD45.2 HSC-derived cells to bone marrow (BM) cellularity relative to their contribution to peripheral blood (*n* = 10). **(J)** Representative histograms of the contribution of CD45.2+ cells (HSC donor derived) and CD45.1+ cells (competitor WBM derived) to the LSK compartment. Error bars represent SEM. * represent outliers.

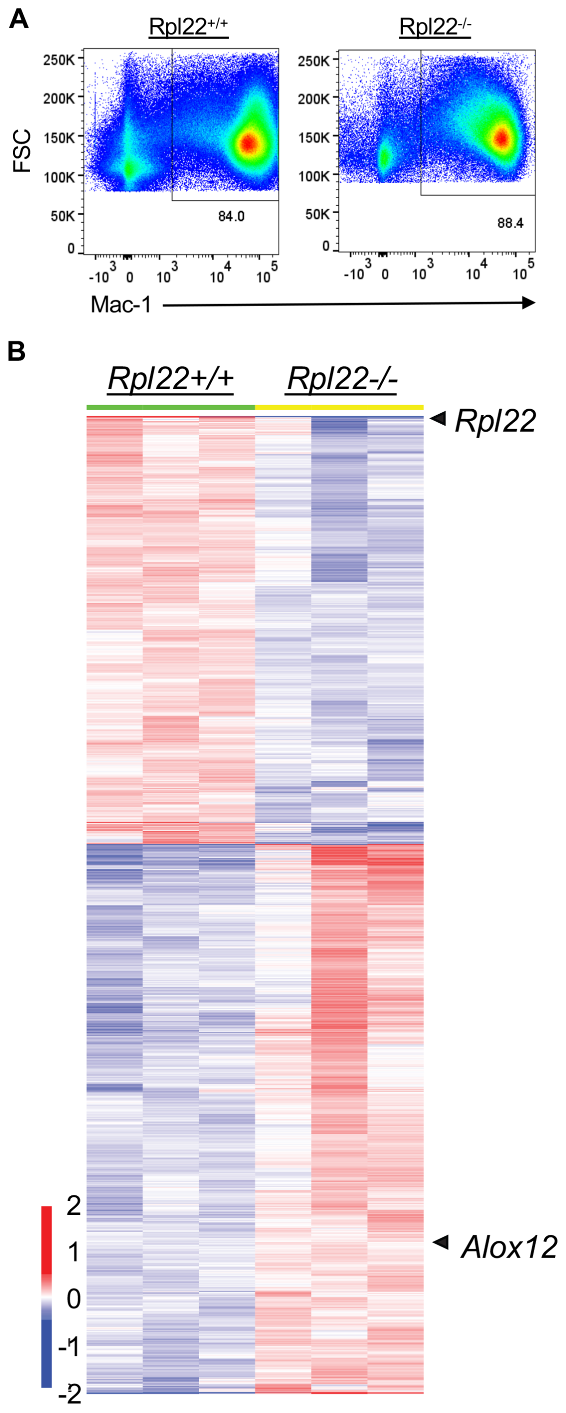


**Figure S3, related to Figure 4: Rpl22-deficiency alters the transcriptomes of adult HSC.** **(A)** Representative flow cytometric analysis of myeloid burden in spleens of MLL-AF9 knockin *Rpl22*$^{+/+}$ and *Rpl22*$^{-/-}$ mice at time of death. Histograms of splenocytes stained with the anti-Mac-1/CD11b, a myeloid marker, are depicted. **(B)** The heat map depicts genes differentially expressed in triplicate samples of HSC isolated from *Rpl22+/+* and *Rpl22-/-* mice. *Rpl22* and *Alox12* are indicated. Significance was defined as $p < 0.05$ and FDR < 20%.

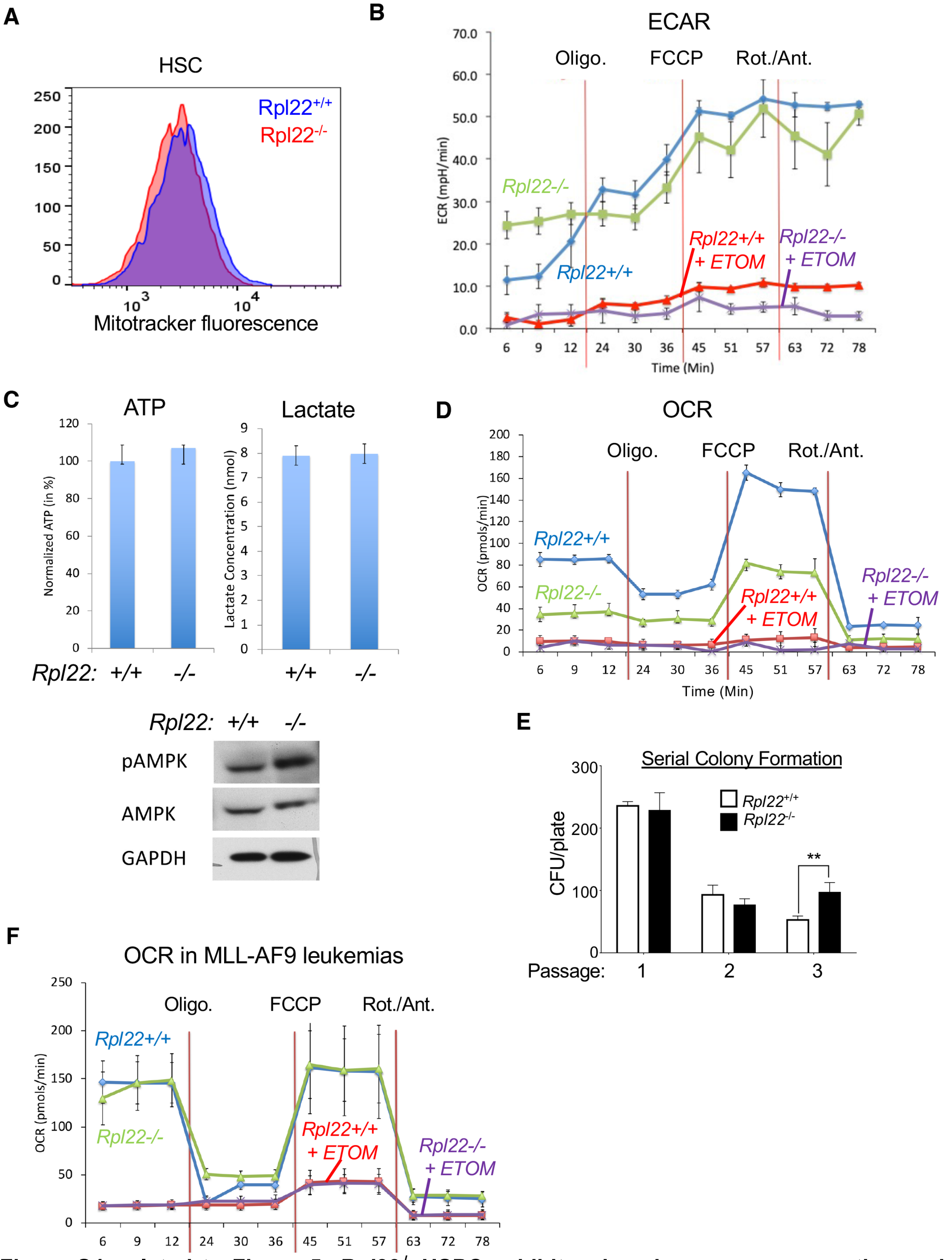


**Figure S4, related to Figure 5: *Rpl22*$^{-/-}$ HSPC exhibit reduced oxygen consumption and sustained colony forming ability. (A)** Representative histogram of measurement of mitochondrial biomass using Mitotracker staining. **(B)** The extracellular acidification rate (ECAR) of *Rpl22*$^{+/+}$ and *Rpl22*$^{-/-}$ LSK, a measure of glycolytic potential, was assessed using the Seahorse XFe96 Bioanalyzer and the Seahorse Cell Energy Phenotyping kit. Triplicate measures were performed with and without treatment with etomoxir, an inhibitor or FAO. The Representative graph of the mean +/- SEM illustrates the timing of addition of oligomycin (Oligo.), FCCP, rotenone (Rot), and antimycin A (Ant.). **(C)** The ATP and lactate content of *Rpl22*$^{+/+}$ and *Rpl22*$^{-/-}$ LSK was measured using colorimetric assay kits from Biovision. The normalized mean +/- SEM of triplicate measures is depicted graphically. The level and phosphorylation of AMPK was assessed by immunoblotting with GAPDH serving as loading control (lower panels). **(D)** The oxygen consumption rate (OCR), an indicator of aerobic respiration, was quantified on *Rpl22*$^{+/+}$ and *Rpl22*$^{-/-}$ LSK using the Seahorse Cell Energy Phenotyping kit. The representative graph depicting the mean +/- SEM of triplicate measures and the timing of addition of the same inhibitors as in **(B)** are shown. **(E)** Graphical representation of colony forming ability of *Rpl22*$^{+/+}$ and *Rpl22*$^{-/-}$ LSK through three serial passages. The mean +/- SEM of triplicate measures of colonies formed per plate is depicted graphically. **(F)** The OCR was quantified on MLL-AF9 transformed *Rpl22*$^{+/+}$ and *Rpl22*$^{-/-}$ leukemias using the Seahorse Cell Energy Phenotyping kit. The representative graph depicting the mean +/- SEM of triplicate measures and the timing of addition of the same inhibitors as in **(D)** is shown. All experiments were performed three times and graphs were analyzed for significance using the student's t-test. Error bars represent SEM. $^{**}p \leq 0.01$.

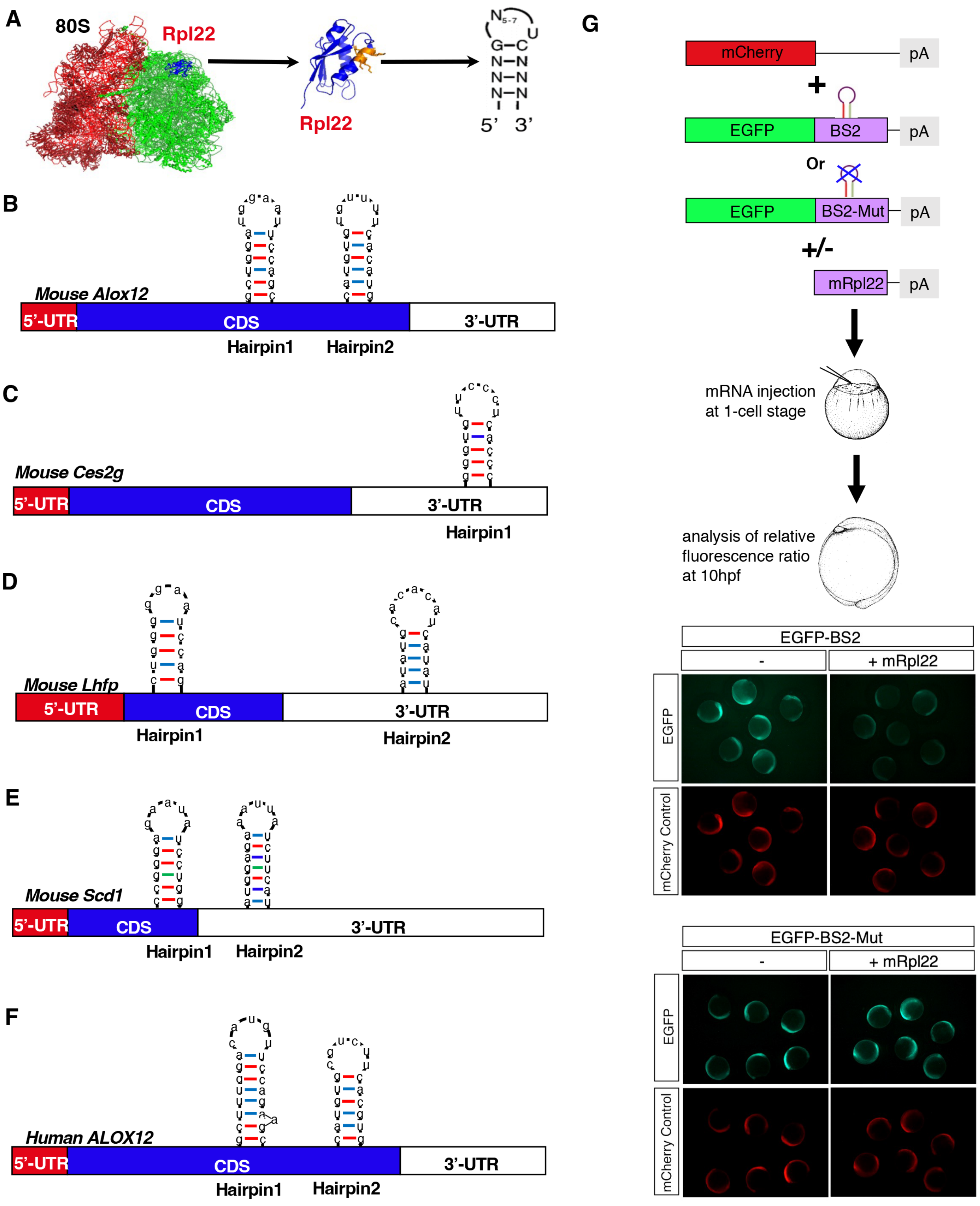


**Figure S5, related to Figure 5: Rpl22 binding motifs in differentially expressed mRNA encoding regulators of FAO. (A)** Schematic illustrating the location of Rpl22 protein in the crystal structure of the 80S ribosome, a ribbon diagram of the crystal structure of Rpl22, and the consensus stem-loop structure bound by Rpl22. **(B-F)** Diagrammatic representation of the location of predicted, consensus Rpl22 stem-loop binding sites in mRNA encoding murine **(B)**

*Alox12,* (**C**) *Ces2g,* (**D**) *Lhfp,* (**E**) *Scd1,* and human *ALOX12* **(F)**. **(G)** The ability of Rpl22 to directly control Alox12 expression was assessed by appending intact or mutated Rpl22 binding sites from BS2 of *Alox12* mRNA onto a GFP reporter. The intact and mutant Alox12-GFP reporter, along with an mCherry injection control, were injected into 1 cell stage zebrafish embryos alone or with mRNA encoding Rpl22. The ability of Rpl22 to bind and repress through the *Alox12* binding site was assessed by quantifying the effect on GFP fluorescence. Representative images are depicted in the bottom panels **(G).**

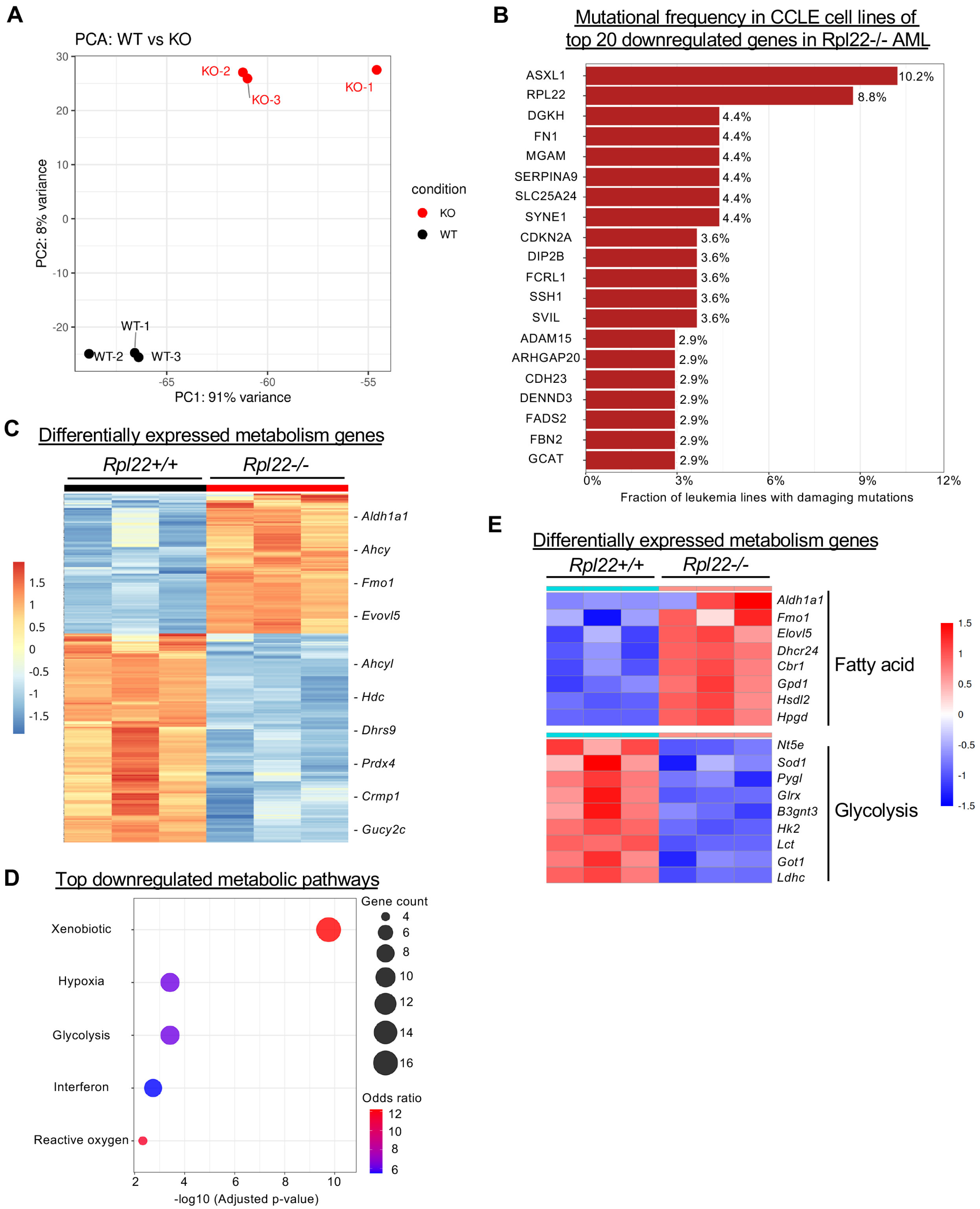


**Figure S6, related to Figure 7: Expression profiling on Rpl22$^{+/+}$ and Rpl22$^{-/-}$ MLL-AF9 leukemias. (A)** Principal component analysis of triplicate samples of explanted Rpl22$^{+/+}$ and Rpl22$^{-/-}$ MLL-AF9 leukemias analyzed by RNA-Seq. WT, Rpl22$^{+/+}$; KO, Rpl22$^{-/-}$ **(B)** Graphical representation of the frequency with which the top 20 genes that are downregulated in Rpl22$^{-/-}$ leukemias are mutationally inactivated in CCLE human leukemia cells lines. **(C)** Heat map displaying 262 genes differentially expressed in Rpl22$^{-/-}$ MLL-AF9 leukemias that are also involved in regulating metabolic processes. **(D)** Bubble plot of the top downregulated Reactome metabolic pathways in Rpl22$^{-/-}$ MLL-AF9 leukemias, with the bubble size indicating the gene count and color indicating p value. **(E)** Heat map of differentially expressed genes in Rpl22$^{-/-}$ MLL-AF9 leukemias involved in fatty acid metabolism and glycolysis.

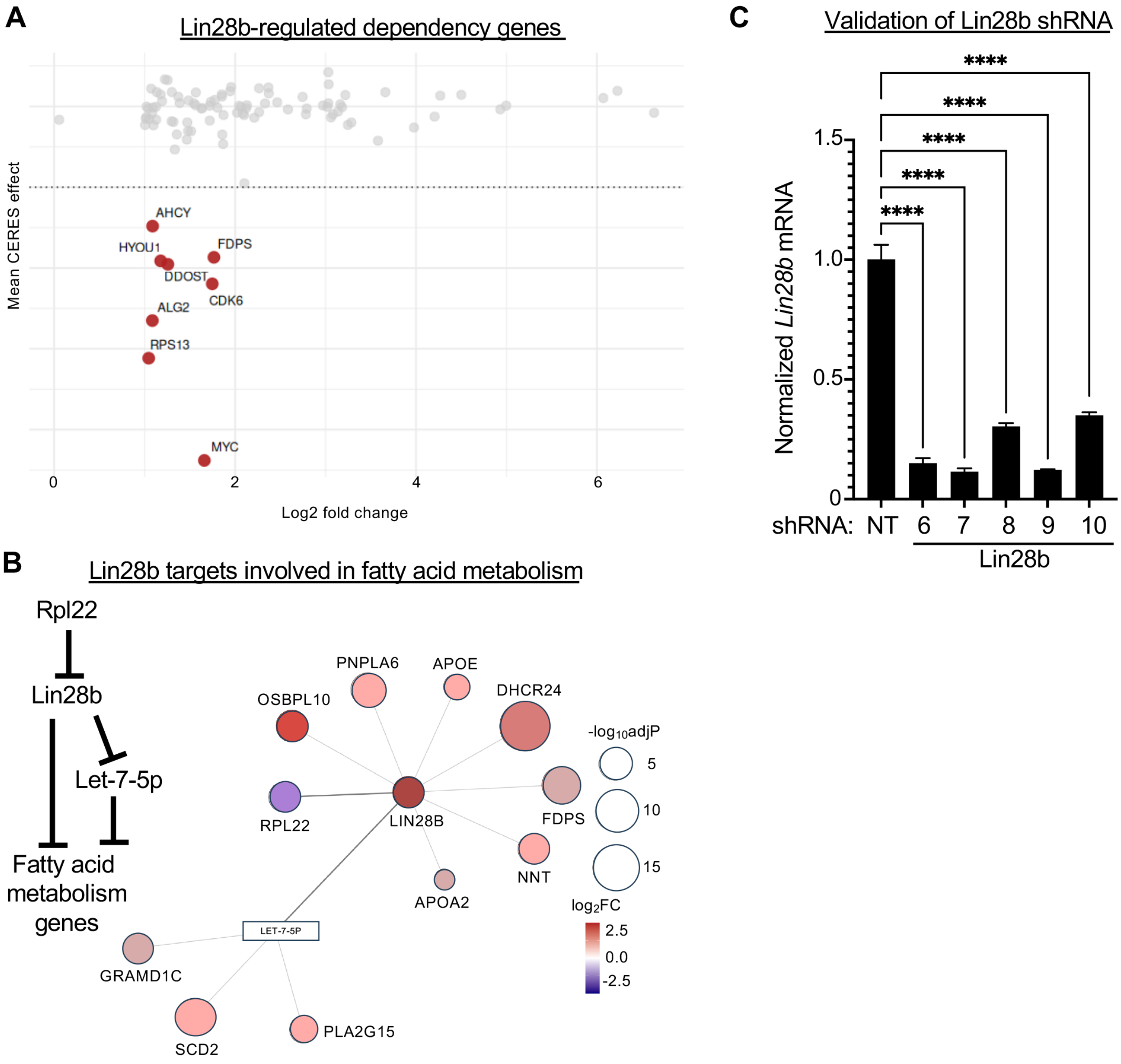


**Figure S7, related to Figure 7: Role of Lin28b in regulating lipid content of Rpl22-/- MLL-AF9 leukemias. (A)** Scatter plot of mean CRISPR dependency score versus Log2 fold-change for differentially expressed Lin28b gene targets induced in Rpl22-/- MLL-AF9 leukemia. **(B)** qPCR measurement of the impact of *Lin28b* shRNA on *Lin28b* mRNA levels, normalized to a non-targeting (NT) control. Statistical significance was determined by one-way ANOVA. **** p<0.0001. **(C)** String network analysis of Lin28b targets induced in Rpl22-/- MLL-AF9 leukemias and in involved regulating lipid metabolism.
