## Supplementary figures and images for "Ribosomal protein control of hematopoietic stem cell transformation through regulation of metabolism"

### Harris et al., 2025 Table S5.tif

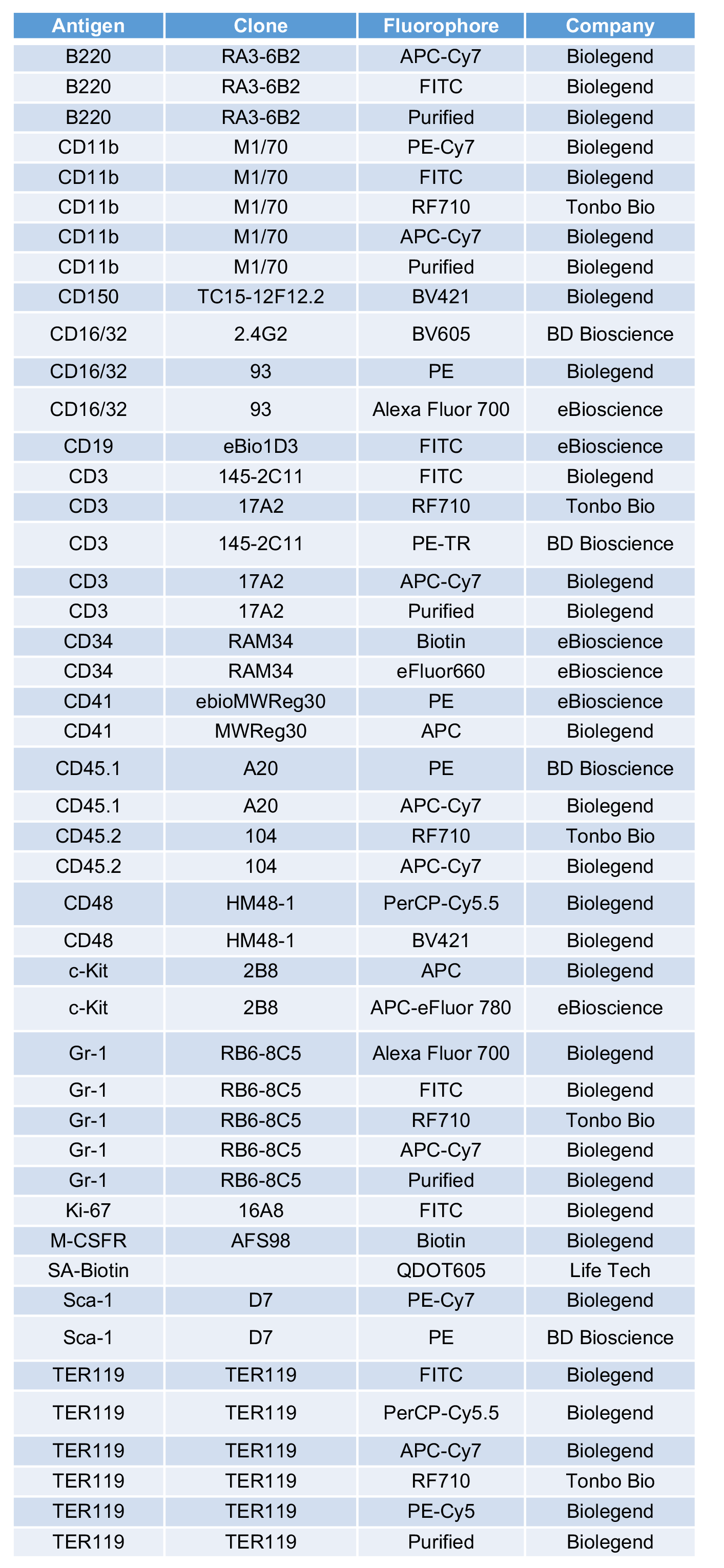
